# HERC4 limits oxidative stress–induced DNA damage during bacterial and viral-bacterial infection

**DOI:** 10.64898/2026.07.02.736007

**Authors:** Clemens Cammann, Vanessa Gering, Thomas Sura, Abhishek K. Singh, Julia D. Boehme, Eylin Topfstedt, Anne K. Koch, Ulrike Ritter, Karsten Becker, Dunja Bruder, Ulrike Blohm, Hortense Slevogt, Sandra Maaß, Gernot Rohde, Jan Rupp, CAPNETZ Study Group, Sven Hammerschmidt, Dörte Becher, Ulrike Seifert

## Abstract

Respiratory (co-)infections caused by influenza viruses and *Streptococcus pneumoniae* represent significant threats to global health. In our analysis of host cell ubiquitination, we identified reactive oxygen species (ROS) produced by *S. pneumoniae* as critical effectors in reducing the amount of intracellular polyubiquitinated proteins. Together with reduced ubiquitination we observed a downregulation of the E3 ligase HERC4 upon infection with *S. pneumoniae* in human alveolar epithelial and macrophage-like cells as well as in samples obtained from *S. pneumoniae* infected humans and mice. This was further aggravated in the viral-bacterial coinfection with influenza A. CRISPR-Cas9 deletion of HERC4 prior bacterial infection resulted in increased ROS-induced DNA damage, enhanced host cell apoptosis and reduced Histone 2B ubiquitination. In contrast, HERC4 overexpression diminished DNA damage indicating a role of HERC4 in DNA-damage-repair upon infection. By establishing a link between HERC4 expression and ROS-induced DNA damage and repair, we identified a potential marker for predicting the outcome of viral and bacterial (co-)infections. Targeting HERC4 expression defines a novel strategy to protect host cells from *S. pneumoniae* (co-)infection attenuating infection exacerbation.

## Introduction

Influenza A virus (IAV) and *Streptococcus pneumoniae* represent two major pathogens of respiratory tract infections and disease. While IAV can cause a clinical spectrum that ranges from asymptomatic infections to respiratory failure and acute respiratory distress syndrome (ARDS) (1), *S. pneumoniae* is the leading pathogen in community-acquired pneumonia (CAP), but often persist as an asymptomatic colonizer of the upper respiratory tract, indicating that infectious disease manifestation depends on the balance between bacterial virulence, host susceptibility and local immune control (2). During influenza-associated co-infection, pneumococci are among the most common bacterial pathogens and can exploit virus-induced alterations in lung epithelial barrier integrity and antibacterial host defenses (3). Viral-bacterial coinfection is associated with increased disease severity and mortality compared with single infections (4). In clinical settings, distinguishing viral from bacterial pneumonia is often difficult, and antibiotic therapy is therefore frequently initiated empirically, contributing to substantial antibiotic misuse (2).

Respiratory infections trigger immune and inflammatory responses that can be associated with excessive oxidative stress that drives cell and tissue damage as well as severe disease outcomes. Oxidative stress is increasingly recognized as a key component of respiratory infection pathogenesis, reflecting an imbalance between infection-induced reactive oxygen species (ROS) production and host antioxidant defense mechanisms (5). In the lung, excessive ROS can promote epithelial injury, barrier dysfunction, DNA damage, as well as alterations in histone modifications and chromatin architecture that shape host transcriptional responses (5, 6). During IAV infection, oxidative stress is driven predominantly by endogenous ROS generated by host cells through mitochondrial respiration and NADPH oxidase–dependent oxidative bursts (7, 8). In contrast, *S. pneumoniae* contributes directly to the local oxidative burden by producing large amounts of hydrogen peroxide via the pyruvate oxidase SpxB which accumulates extracellularly and further promotes oxidative stress, DNA damage, and epigenetic remodeling (9, 10).

Ubiquitination is a highly versatile post-translational modification of eukaryotic proteins that regulates protein turnover, inflammatory signaling, endocytosis, DNA damage responses, and host–pathogen interactions (11). It occurs either as mono-ubiquitination, involving the covalent attachment of a single ubiquitin moiety to a substrate, which primarily influences protein localization, trafficking, and functional activity, or as poly-ubiquitination, in which ubiquitin chains are assembled through linkage to one of the seven lysine residues or the N-terminal methionine of ubiquitin itself (12). The biological consequences of poly-ubiquitination are determined by the specific chain topology: for instance, K48-linked ubiquitin chains predominantly target proteins for proteasomal degradation, whereas K63-linked chains primarily mediate non-degradative functions, including innate immune signaling, receptor trafficking, and DNA damage repair. The specificity of ubiquitin signaling is largely conferred by E3 ubiquitin ligases, which recognize defined substrates and determine the timing, localization, and architecture of ubiquitin modifications, thereby directing distinct cellular outcomes. Among E3 ubiquitin ligases, homologs to the E6-associated protein C-terminus (HECT)-type ligases are important for their roles in stress-responsive signaling pathways that intersect with cellular redox homeostasis, DNA damage responses, and protein quality control (13). Within this context, HERC4 belongs to the HECT and regulator of chromatin condensation 1-like domain-containing protein (HERC) subfamily of E3 ligases and has been implicated in diverse cellular pathways, including cell migration, apoptosis, and viral infection (14). Depending on the substrate, HERC4 has so far been described to mediate K48-linked and atypical K63-linked poly-ubiquitination (15, 16). It is ubiquitously expressed across tissues, with particularly high expression levels in the brain and testis (17), and is predominantly localized to the cytoplasm, although it has also been observed in the nucleus (18). Its physiological function has mainly been associated with spermatogenesis, however, dysregulated HERC4 expression has also been reported in several cancers, where it may correlate with disease stage and has therefore been proposed as a potential biomarker (19, 20). To date, however, no studies have investigated the susceptibility of HERC4 to oxidative stress or its role during pathogen infection.

In infection, ubiquitination is required for multiple antiviral and antibacterial host responses, but pathogens can also exploit ubiquitin-dependent pathways to promote replication, persistence, or immune evasion. In IAV infection, ubiquitination contributes to viral entry, endocytosis, replication, and the proteasome-dependent turnover of viral and host proteins (21–23). In bacterial infection, ubiquitin-dependent host defense has been extensively characterized in intracellular infection models, in which cytosolic bacteria become decorated with ubiquitin chains that recruit selective autophagy receptors and activate inflammatory signaling pathways (24–26). Although selected extracellular pathogens have been shown to engage ubiquitin-dependent host pathways through toxins, translocated effectors, or specific host substrates, a systematic understanding of how extracellular respiratory bacteria reshape host cell ubiquitination remains limited.

In this context, *S. pneumoniae* represents a clinically relevant extracellular respiratory pathogen. Pneumococcal infection is linked to oxidative stress, DNA damage, severe pneumonia, and influenza-associated bacterial coinfection. The production of hydrogen peroxide by pneumococci provides a plausible mechanism by which bacterial infection may alter ubiquitin-dependent host cell processes. However, how *S. pneumoniae*-induced oxidative stress intersects with ubiquitin-dependent host cell regulation during bacterial mono- and viral-bacterial coinfection remains incompletely understood.

In this study, we investigated how single infections with *S. pneumoniae* or IAV, as well as IAV-*S. pneumoniae* coinfection, affect host cell ubiquitination. Global ubiquitination patterns were assessed in human alveolar epithelial cells, whereas subsequent analyses focused on the infection-associated regulation of the E3 ligase HERC4 across human epithelial and macrophage-like infection models, mouse-derived alveolar epithelial cells and patient-derived blood samples. We further examined whether bacterial H_2_O_2_ contributes to *S. pneumoniae*-induced modulation of host ubiquitination pathways and whether HERC4 may link bacterial H_2_O_2_-associated DNA damage to host DNA damage response signaling.

## Results

### Reduced levels of poly-ubiquitinated protein conjugates in alveolar epithelial cells during bacterial infection

The ubiquitin-conjugating system is required for viral replication (27–29) and virus infection is accompanied by an increase in the total amount of poly-ubiquitinated proteins in host cells (30). We confirmed these findings by monitoring the overall poly-ubiquitination up to 24h upon influenza A (H1N1)pdm09 (IAV) infection in A549 lung epithelial cells, which displayed a pronounced increase of ubiquitinated protein conjugates (Figure 1A and B and Figure S1A, infection control). In contrast, infection with the Gram-positive bacterium *S. pneumoniae* D39Δ*cps* (serotype 2), a capsule-deficient strain used as a model to foster host–pneumococcal interaction (31, 32), resulted in a significant decrease in poly-ubiquitinated proteins up to 10 h post infection (Figure 1C and D and Figure S1B, infection control).

**Figure 1.**
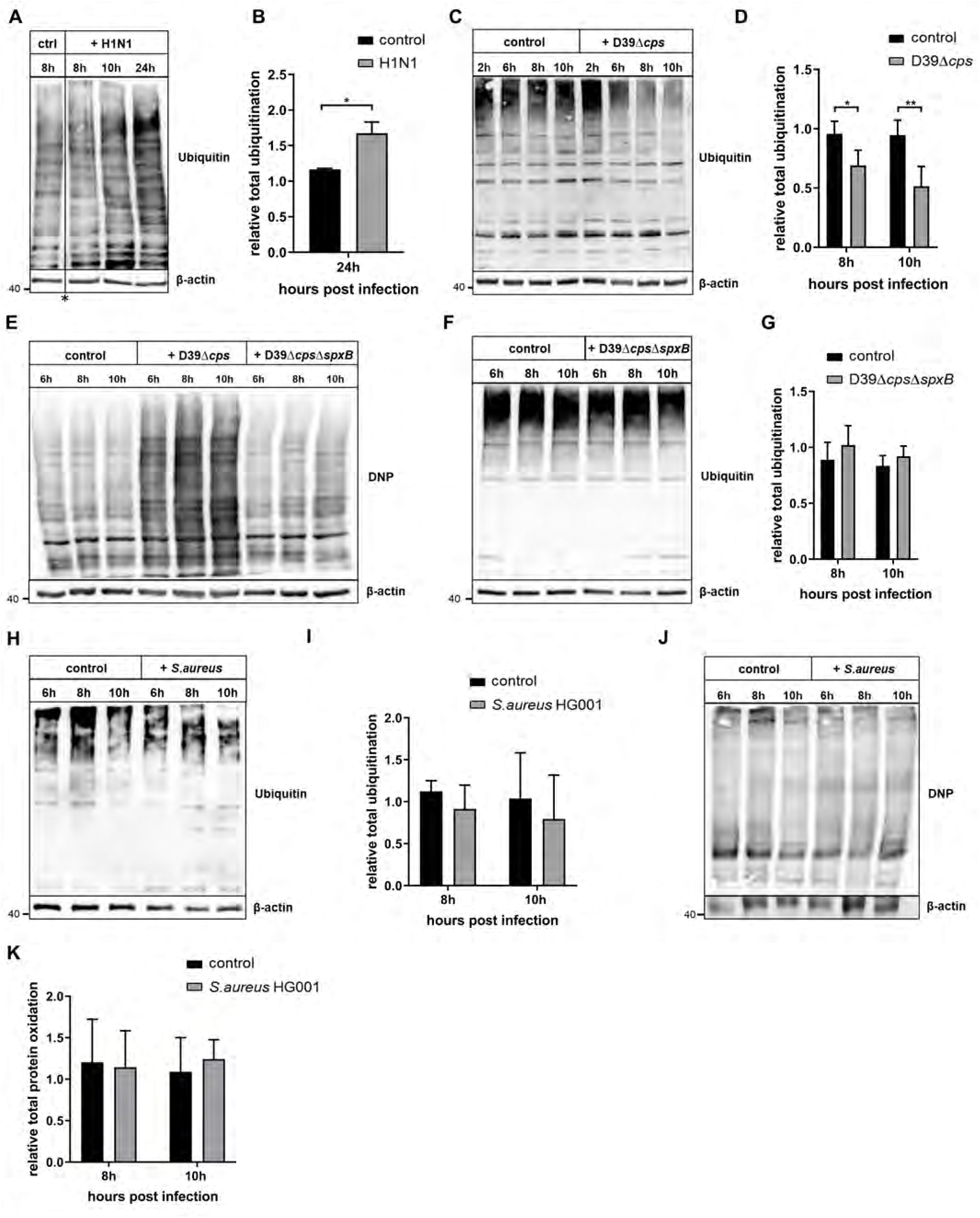
Reduction of total ubiquitination upon *S. pneumoniae*-induced oxidative stress in human alveolar epithelial cells. A, B A549 cells were either infected with influenza A virus (H1N1)pdm09 (MOI 5) or non-infected (ctrl). Cells were harvested at depicted time points and total ubiquitination was assessed by immunoblotting with β-actin as loading control (A). Lane intensities were analyzed by densitometry and normalized to β-actin. Graph depicts relative change in total ubiquitination calculated to the 8 h uninfected control (B), n = 2. * black line indicates cutting of the original blot (A) C, D A549 cells were either infected with *S. pneumoniae* D39Δ*cps* (MOI 20) or non-infected (control). Cells were harvested at depicted time points and total ubiquitination was assessed by immunoblotting with β-actin as loading control (C). Lane intensities were analyzed by densitometry and normalized to β-actin. Graph depicts relative change in total ubiquitination calculated to 6 h uninfected control (D) n = 3. E A549 cells were analyzed for the amount of oxidized proteins via OxyBlot upon infection with *S. pneumoniae* D39Δ*cps* (MOI 20) and *S. pneumoniae* D39Δ*cps*Δ*spxB* (MOI 20) or left untreated (control) using anti-(2,4-dinitrophenol) DNP antibodies against derivatized carbonyl groups in oxidized proteins, n = 3. F, G A549 cells were infected with *S. pneumoniae* D39Δ*cps*Δ*spxB* (MOI 20) or left untreated (control). Cells were harvested at depicted time points and total ubiquitination was assessed by immunoblotting with β-actin as loading control (F). Lane intensities were analyzed by densitometry and normalized to β-actin. Graph depicts relative change in total ubiquitination calculated to respective 6 h time point (G), n = 3. H, I A549 cells were either infected with *S. aureus* HG001 (MOI 20) or non-infected (control). Cells were harvested at depicted time points and total ubiquitination was assessed by immunoblotting with β-actin as loading control (H). Lane intensities were analyzed by densitometry and normalized to β-actin. Graph depicts relative change in total ubiquitination calculated to 6 h uninfected control (I), n = 3. J, K A549 cells were analyzed for the amount of oxidized proteins via OxyBlot upon infection with *S. aureus* HG001 (MOI 20) or left untreated (control) by analyzing the immunoblot with anti-DNP antibodies against derivatized carbonyl groups in oxidized proteins (J). Lane intensities were analyzed by densitometry and normalized to β-actin. Graph depicts relative change in total protein oxidation calculated to respective 6 h time point (K), n = 3. Data information: in B, D, G, I and K data are presented as mean ± SD. \**p* < 0.05; \*\**p* < 0.01. (student’s t test)

Extracellular reactive oxygen species (ROS) such as hydrogen peroxide (H_2_O_2_) derived from *S. pneumoniae* are prominent virulence factors contributing to the severity of infection (33). High amounts of ROS are responsible for host cell protein oxidation which in turn leads to the rapid degradation of oxidized proteins by either the proteasome or lysosomal pathways (34–37). To further explore the relationship between *S. pneumoniae-*induced oxidative stress and changes in the host protein ubiquitination machinery, we compared infections with the *S. pneumoniae* strain D39Δ*cps* and D39Δ*cps*Δ*spxB*, which lacks the pyruvate oxidase and therefore exhibits markedly reduced H_2_O_2_ production (38). As expected and shown before (39), infection with the *S. pneumoniae* D39Δ*cps* strain led to a considerable accumulation of oxidized proteins as evidenced by the enhanced detection of carbonylated proteins. In contrast, non-treated A549 control cells and cells infected with the D39Δ*cps*Δ*spxB* mutant showed no increase in protein oxidation (Figure 1E and Figure S1C, infection control). In addition, infections with D39Δ*cps*Δ*spxB* largely restored poly-ubiquitination (Figure 1F and G). These data indicate that the decrease in poly-ubiquitinated conjugates during *S. pneumoniae* D39Δ*cps* infection coincides with an increase of oxidant-damaged host cell proteins due to ROS release by the bacteria.

To assess whether H_2_O_2_ contributes to the observed effects, we performed infection experiments with *Staphylococcus aureus*, a Gram-positive bacterium that generates only minor amounts of H_2_O_2_ and efficiently detoxifies reactive oxygen species derived from bacteria or host by expressing catalase (40). In contrast to *S. pneumoniae* D39Δ*cps*, infection with *S. aureus* HG001 revealed only minor alterations of poly-ubiquitinated conjugates and protein carbonylation in A549 alveolar epithelial cells (Figure 1H – K and Figure S1D, infection control). These effects were markedly weaker compared to infection with *S. pneumoniae* D39Δ*cps*, indicating only a modest and non-significant induction of protein modification upon *S. aureus* infection.

Together, these findings suggest a link between bacteria-derived ROS (H_2_O_2_), oxidative stress-induced host cell protein carbonylation, and changes in the abundance of poly-ubiquitinated proteins in *S. pneumoniae* infected cells.

### Altered expression of ubiquitinating enzymes during infection with *S. pneumoniae*

The transfer of ubiquitin moieties to target protein substrates by E3 ligases represents a critical regulatory step in the ubiquitination cascade. To gain molecular insights into the alterations of the host cell ubiquitination machinery during infection, we systematically assessed the abundance of E3 ligases in IAV and *S. pneumoniae* D39Δ*cps*-infected A549 cells using a mass spectrometry-based proteomic approach. Analysis of published proteomic data from IAV-infected A549 cells published by Sura et al. (41) revealed 70 E3 ligases, of which 4 displayed increased abundance, while no E3 ligase showed reduced expression levels (Figure 2A). In contrast, pneumococcal infections were associated with a distinct pattern. Among the 90 E3 ligases detected, 11 were downregulated, whereas only 1 was upregulated upon *S. pneumoniae* D39Δ*cps* infection (Figure 2B and S1E, infection control). Consistent with this observation, we additionally detected 3 E2 ubiquitin-conjugating enzymes with reduced protein expression levels (Figure EV1A). These results demonstrate that pneumococcal infections are accompanied by a reduced abundance across several components of the host ubiquitination machinery.

**Figure 2.**
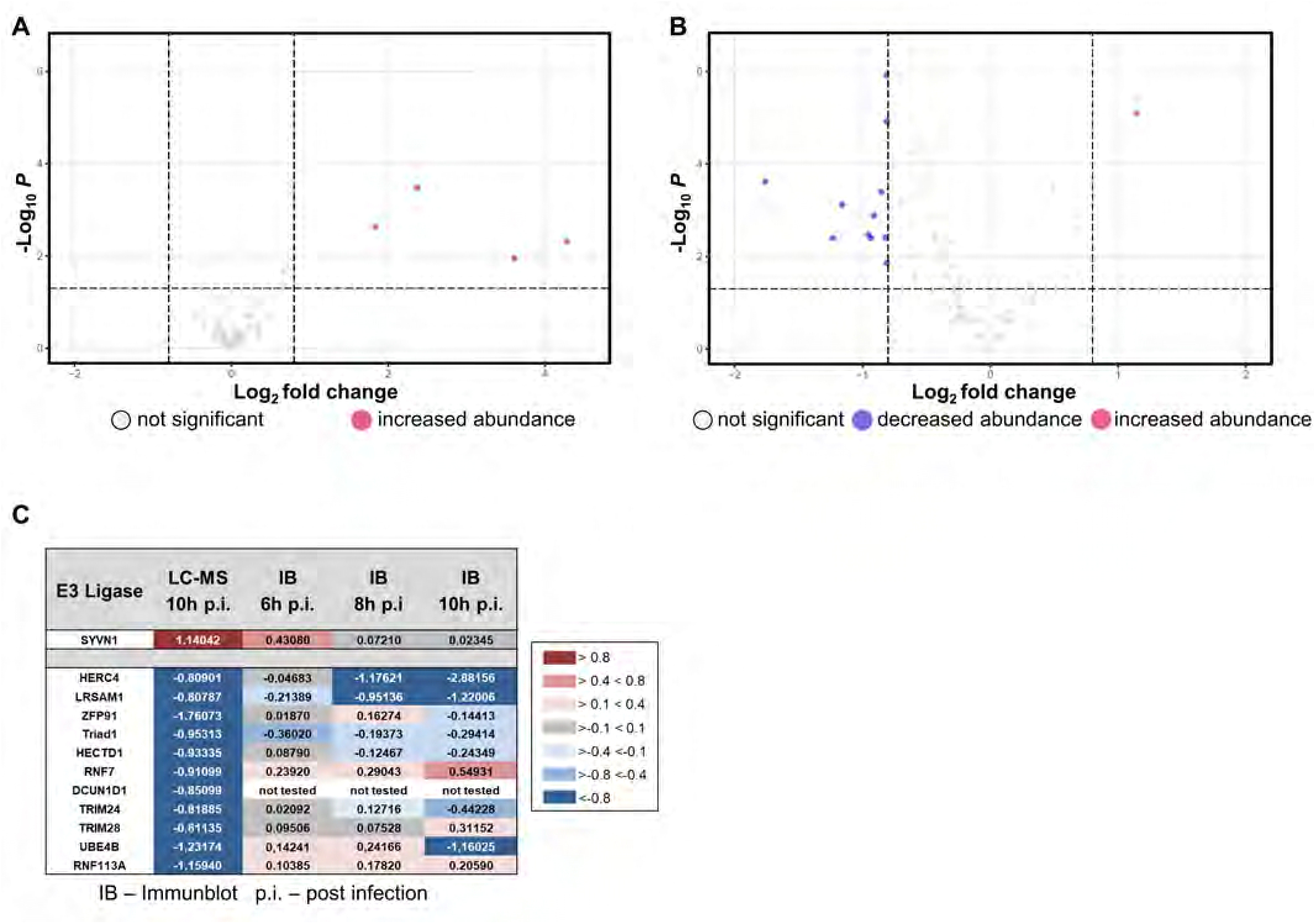
Reduced abundance of E3 ligases during S. pneumoniae infection. A A549 cells were infected with influenza A virus (H1N1)pdm09 (MOI 5) and were harvested after 24 h and subsequently analyzed by LC-MS for E3 ligase abundance, n = 4. Volcano plot of all quantified E3 Ligases generated from previously published proteomics data (41). In total 70 E3 ligases were identified from which 4 showed increased abundance (red dots) and 66 remained unchanged (grey dots). Proteins were considered as differentially abundant with a fold change > 1.74 indicated as log2foldchange > |0.8|, with p < 0.05. B A549 cells were infected with *S. pneumoniae* D39Δ*cps* (MOI 15) and harvested after 10 h and subsequently were analyzed by LC-MS for E3 ligase abundance, n = 4. In total 89 E3 ligases were identified of which 10 showed decreased abundance (blue dots), 1 increased abundance (red dots), and 78 remained unchanged (grey dots). Proteins were considered as differentially abundant with a fold change > 1.74 indicated as log2foldchange > |0.8|, with p < 0.05. C Comparison of differentially expressed E3 ligases upon infection with S*. pneumoniae* D39Δ*cps* (MOI 20) which showed a significant increased (red) or decreased (blue) abundance in the LC-MS analysis after 10 h infection (n = 4) with the respective immunoblot (IB) results after 6 h, 8 h, and 10 h infection (n = 2), values are calculated to the respective uninfected controls as log2fold change.

To further characterize the expression pattern of the E3 ligases identified in Figure 2B, we performed infection kinetics and analyzed the protein expression by immunoblotting (Figure 2C and Figure EV1B and S1F, infection control). While some E3 ligases, in particular RNF7, displayed expression patterns that diverged from the Liquid Chromatography-Mass Spectrometry (LC-MS) data, the majority of the analyzed E3 ligases showed concordant results, characterized by a downregulation 10 h post infection (Figure 2C and Figure EV1B). For HERC4 and LRSAM1 kinetic analysis revealed sustained downregulation throughout the infection time course in A549 cells (Figures 2C, 3A – C and D – E, infection controls). In case of HERC4 downregulation could be confirmed in primary small airway epithelial cells (SAEC), already at 6 h and 8 h after infection (Figure EV2A – B and C – D, infection controls).

**Figure 3.**
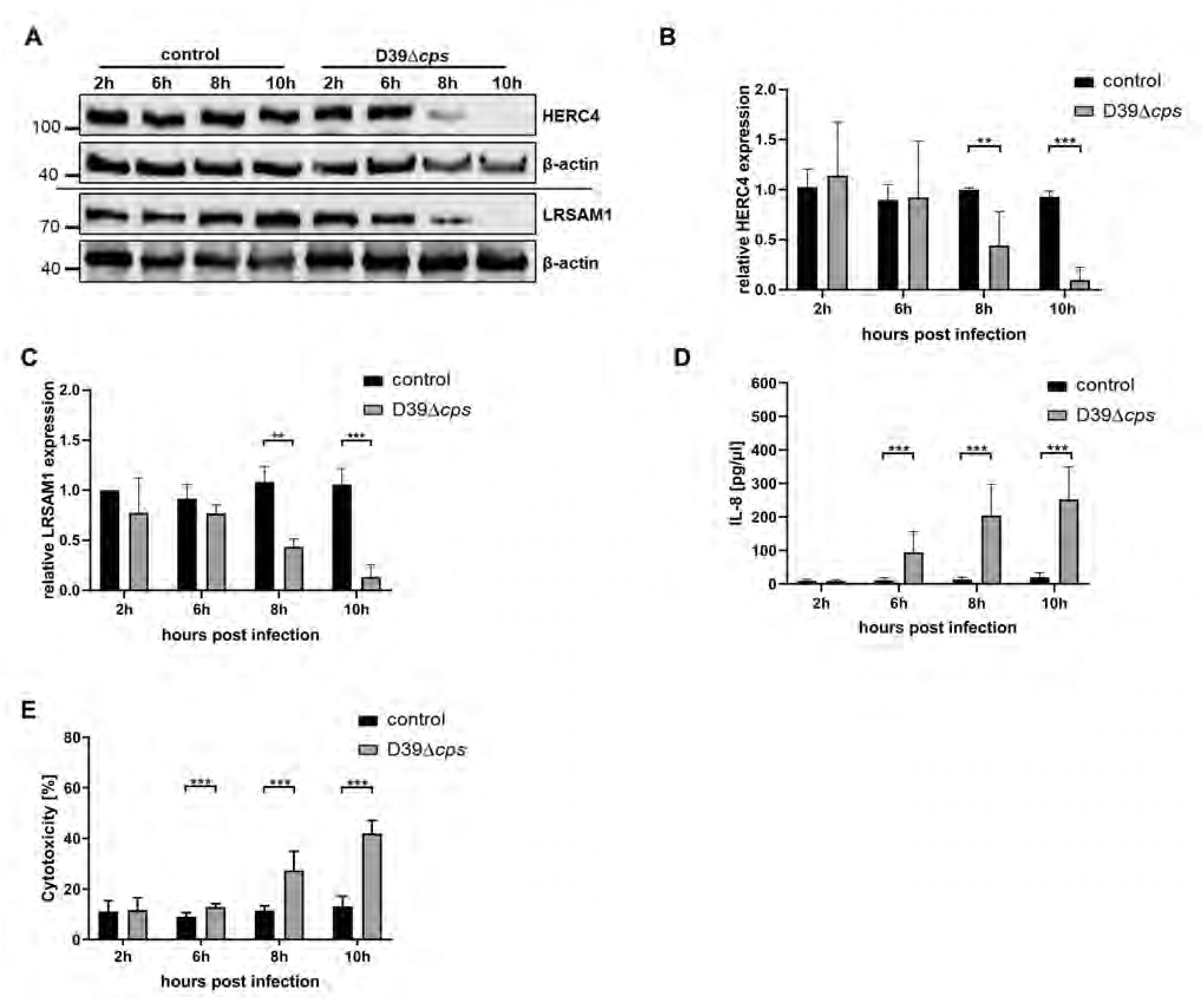
Decreased HERC4 and LRSAM1 expression during infection with S. pneumoniae D39Δ*cps* in human alveolar epithelial cells. A-E A549 cells were infected with *S. pneumoniae* D39Δ*cps* (MOI 20) for the depicted time points and subsequently analyzed for HERC4 and LRSAM1 protein expression by immunoblotting with β-actin as loading control (A). Band intensities were analyzed by densitometry and normalized to β-actin. Graphs depict relative expression of HERC4 (B) and LRSAM1 (C) calculated to 2 h uninfected control, n = 3. Infection controls: cell culture supernatants were analyzed for secretion of IL-8 by ELISA (D), n = 3, and cytotoxicity was monitored by determining extracellular lactate dehydrogenase (E), n = 3. Data information: in B-E data are presented as mean ± SD. \*\**p* < 0.01; \*\*\**p* < 0.001. (student’s t test)

### Pneumococcal-induced oxidative stress regulates HERC4 stability during infection

To investigate the upstream signaling events contributing to the downregulation of HERC4 and LRSAM1 in infected host cells, we analyzed the involvement of the cell surface pattern recognition receptors Toll-like receptor 2 (TLR2) and Toll-like receptor 4 (TLR4) known to mediate recognition of *S. pneumoniae* by host cells during infection (42, 43). TLR2 and TLR4 inhibition prevented the degradation of LRSAM1, however inhibition of either TLR2 or TLR4 failed to rescue HERC4 levels in *S. pneumoniae-*infected alveolar epithelial cells (Figure 4A and Figure S2A and B, infection control).

**Figure 4.**
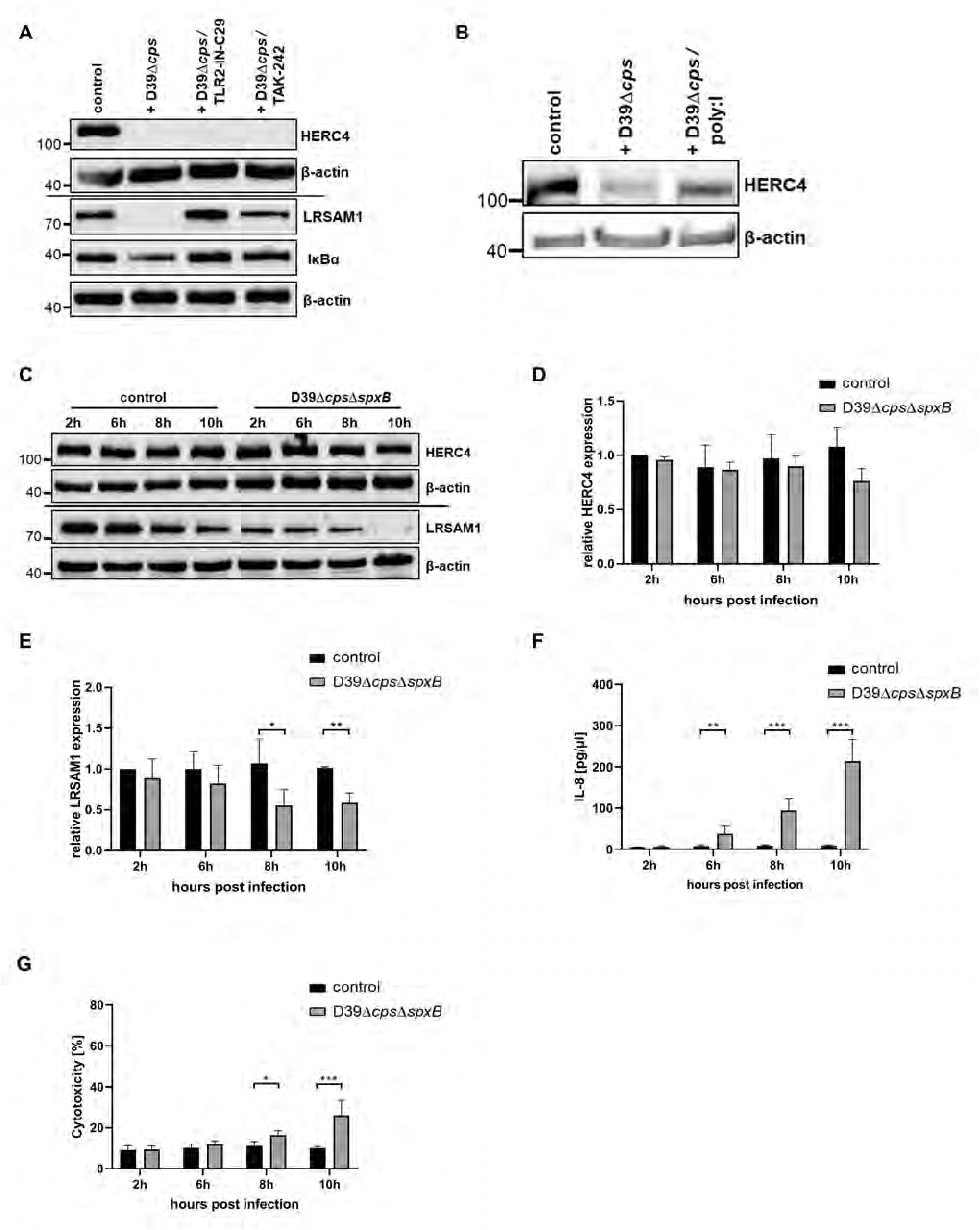
Stable HERC4 expression upon infection with *S. pneumoniae* D39Δ*cps*Δ*spxB* in A549 cells. A, B A549 cells were analyzed for HERC4, LRSAM1, and IκBα protein expression (control for inhibition of NF-kB signaling in the presence of TLR inhibitors) by immunoblotting 8 h post infection with *S. pneumoniae* D39Δ*cps* (MOI 20) in the presence of TLR2 inhibitor TLR2-IN-C29 (200 µM), TLR4 inhibitor TAK-242 (5µM) (A), n = 2, and scavenger receptor inhibitor poly:I (20µg / mL) (B), n = 2, compared to untreated infected and non-infected cells (control), β-actin loading control. C–G A549 cells were analyzed for HERC4 and LRSAM1 protein expression at the depicted time-points during infection with *S. pneumoniae* D39Δ*cps*Δ*spxB* (MOI 20) and compared to non-infected controls (C). Band intensities were analyzed by densitometry and normalized to the loading control β-actin. Graph depicts relative expression of HERC4 (D) and LRSAM1 (E) compared to 2 h uninfected control, n = 4. Infection controls: secretion of IL-8 was determined during infection in the cell culture supernatants by ELISA (F), n = 4 and cytotoxicity was monitored by determining extracellular lactate dehydrogenase (G), n = 4. Data information: in D - G data are presented as mean ± SD. \**p* < 0.05; \*\**p* < 0.01; \*\*\**p* < 0.001. (student’s t test)

These findings raised the possibility that scavenger receptors (SRs), another class of pattern recognition receptors involved in the innate immune response to *S. pneumoniae*, contribute to the regulation of HERC4 protein levels (44). Exposure of alveolar epithelial cells to the pan-SR inhibitor poly:I during pneumococcal infection resulted in stabilization of HERC4 (Figure 4B and S2A and B, infection control), suggesting that scavenger receptor activation by *S. pneumoniae* cell wall components such as lipoteichoic acid and/or peptidoglycan contributes to the degradation of HERC4 upon infection (45–49). Based on our observation that protein carbonylation and poly-ubiquitinated protein conjugate formation largely depend on *S. pneumoniae-*derived hydrogen peroxide (Figure 1E and F) we next examined whether HERC4 levels are also modulated by H_2_O_2_. Infections with D39Δ*cps*Δ*spxB* supported this hypothesis since reduction of *S. pneumoniae-* released H_2_O_2_ resulted in stable HERC4 expression whereas LRSAM1 was degraded (Figure 4C – E and F - G, infection control). Accordingly, treatment of A549 cells with H_2_O_2_ resulted in a significant reduction in HERC4 abundance both on protein and mRNA level, similar to that observed upon *S. pneumoniae* D39Δ*cps* infection (EV3A - D and 3A - B).

In contrast to epithelial cells, macrophages are professional phagocytes adapted to tolerate the high ROS-levels they generate as part of their antimicrobial defense system during infection. We therefore asked whether HERC4 expression is similarly altered in macrophages. Infection of macrophage-like THP-1 cells resulted in a pronounced downregulation of HERC4 on protein-level, already at 2 h post infection (Figure EV3E - F). To monitor THP-1 infection we assessed inflammasome activation analyzing IL1β generation via cleavage of pro-IL1β and cellular damage (Figure EV3E and G).

The results from our cellular analyses could be confirmed *in vivo*/*ex vivo* analyzing HERC4 protein expression in type II alveolar epithelial cells isolated from the lungs of C57BL/6JOlaHsd mice infected either with *S. pneumoniae* serotype 4 (TIGR4), the clinical isolate serotype 19F or serotype 7F. Infection with TIGR4 known as a strong inducer of oxidative stress (50) and 19F resulted in substantial downregulation of HERC4, whereas HERC4 levels remained stable upon infection with 7F pointing to a role of bacterial ROS in the reduction of HERC4 in host cells also *in vivo* (Figure EV3H and I, infection control).

To determine the intracellular fate of the HERC4 protein we treated A549 cells with the proteasome inhibitors bortezomib, MG132 or the lysosomal inhibitor bafilomycin during infection. Whereas bortezomib had no effect on HERC4 expression, adding bafilomycin and to a lesser extent MG132 resulted in a stabilization of HERC4 (Figure EV3J).

Thus, our data strongly suggest that *S. pneumoniae*-derived ROS and scavenger receptor signaling regulate HERC4 protein stability. Within host cells HERC4 levels are affected both by infection-mediated reduction of HERC4 transcripts and by proteasome- and/or lysosome-dependent degradation.

### HERC4 in viral, bacterial and viral-bacterial (co-)infection

To investigate whether viral infection modulates HERC4 expression, A549 alveolar epithelial cells were infected with IAV. No changes in HERC4 protein abundance or mRNA levels were detected, consistent with our previous mass spectrometry results (Figure 2A, 5A-C). In line with published data, IAV infection did not increase host cell cytotoxicity, reflecting the reduced replicative fitness and pointing to an attenuated cytopathic effect of H1N1pdm09 compared with pre-2009 seasonal H1N1 strains (Figure S3A - B; (51)).

**Figure 5.**
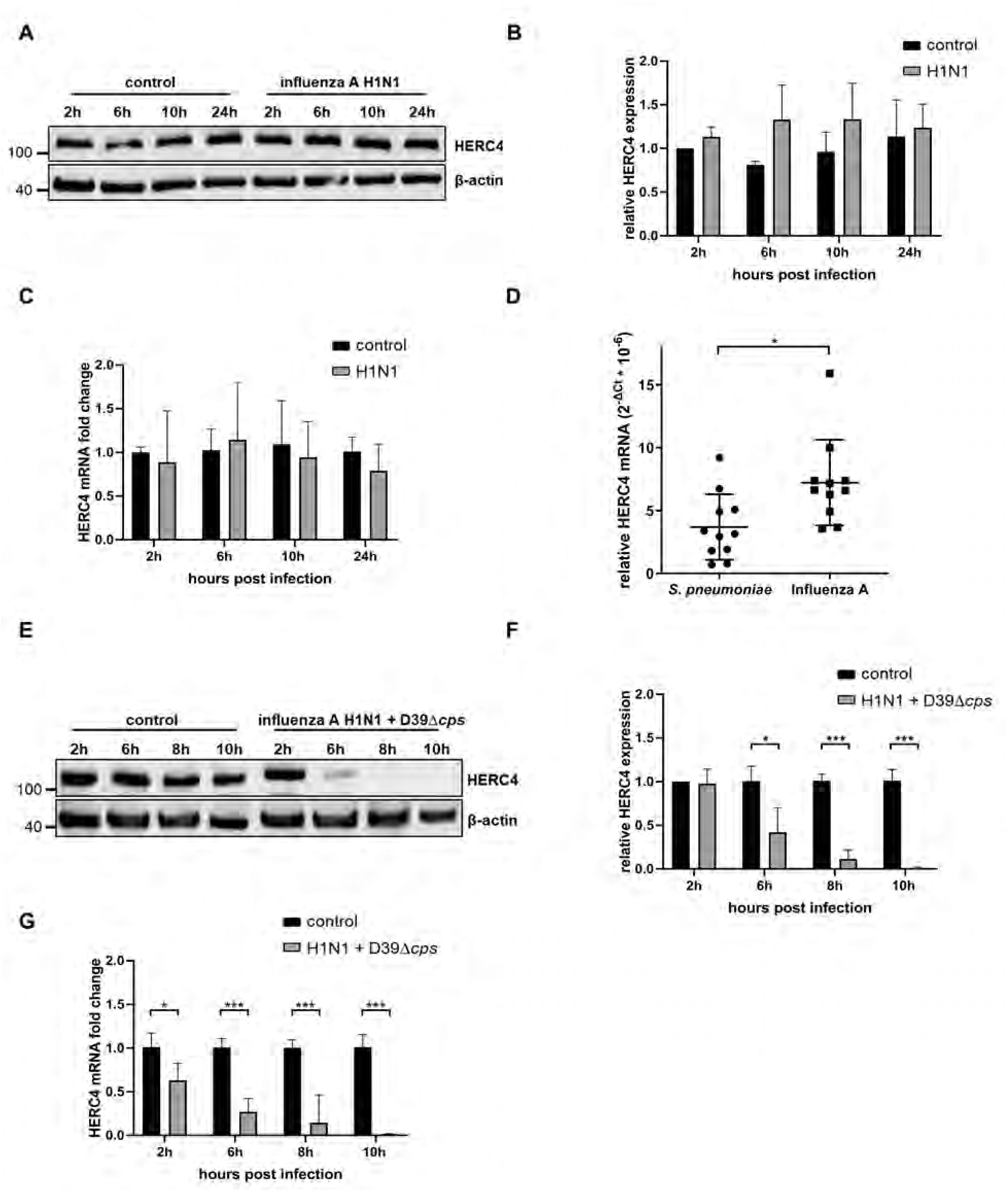
HERC4 down regulation in S. pneumoniae infected patients and in viral-bacterial infected A549 cells. A–C A549 cells were analyzed for HERC4 protein expression at the depicted time points upon infection with influenza A virus (H1N1)pdm09 (MOI 5) or left untreated (control) (A), n = 2. HERC4 band intensities were analyzed by densitometry and normalized to the loading control β-actin. Graphs depict relative expression of HERC4 calculated to 2 h uninfected control (B), n = 2. Changes in HERC4 mRNA were analyzed by qPCR, the fold change was calculated to 2 h uninfected control and normalized to the 18S rRNA housekeeping gene (C), n = 2. D Blood samples collected from patients infected with either *S. pneumoniae* (n = 11) or influenza A virus (n = 11) were analyzed for relative HERC4 expression by qPCR, 2-^ΔCt^ values were calculated for each sample normalized to the 18S rRNA housekeeping gene. E–G A549 cells were analyzed for HERC4 protein expression at the depicted time points upon co-infection with influenza A virus (H1N1)pdm09 (MOI 5) for 24 h followed by bacterial infection with *S. pneumoniae* D39Δ*cps* (MOI 20) (E). HERC4 band intensities were analyzed by densitometry and normalized to the loading control β-actin. Graph depicts relative HERC4 expression calculated to 2 h uninfected control (F), n = 4. Changes in HERC4 mRNA were analyzed by qPCR, the fold change was calculated to the 2 h uninfected control and normalized to the 18S rRNA housekeeping gene (G), n = 4. Data information: in B-D, F and G data are presented as mean ± SD. \**p* < 0.05; \*\*\**p* < 0.001. (student’s t test)

To assess the clinical relevance of our findings, we analyzed HERC4 mRNA expression levels in blood samples from 11 *S. pneumoniae-*infected patients and compared this to 11 blood samples from patients infected with IAV included in the CAPNETZ cohort (52). Influenza A virus infection was confirmed by molecular diagnostics of respiratory samples (including detection of H1N1pdm09), whereas pneumococcal infections were verified in blood cultures. Regarding clinical parameters, lung infiltrates were observed in all patients while fever, cough, elevated CRP levels, and increased leukocyte counts were present in the majority (appendix table 1). A significant lower expression of HERC4 mRNA was detected in patients infected with *S. pneumoniae* compared to IAV-infected individuals (Figure 5D). Taken together, HERC4 downregulation observed in cell experiments and a murine infection model is also evident in human *S. pneumoniae* infection.

IAV infections are often accompanied by secondary coinfections with *S. pneumoniae* which are characterized by a stronger cellular response and higher mortality compared to viral or bacterial mono-infection (53–55). To determine HERC4 expression in viral-bacterial co-infection we infected A549 cells with IAV for 24 h followed by infection with *S. pneumoniae* D39Δ*cps* up to 10 h. Compared to the bacterial mono-infection, we observed an enhanced downregulation of HERC4 on both the protein- and mRNA-level (Figure 5E - G) and a pronounced increase in cell death, confirming the enhanced cytotoxicity associated with co-infection (Figure S3C – E, infection control).

### Deletion of HERC4 enhances bacterial ROS-mediated DNA damage and apoptosis in host cells

Infection-induced ROS can trigger DNA double-strand breaks and can activate the DNA damage response in host cells. Phosphorylation of histone H2AX, a hallmark of this response, has previously been described during pneumococcal infections (50). Consistent with these findings, infection with *S. pneumoniae* D39Δ*cps* significantly increased pH2AX levels (Figure 6A and B and S4A and B, infection control). Phosphorylation of H2AX is mediated by the ataxia-telangiectasia mutated (ATM) protein kinase. Recruitment of the ATM kinase to sites of damaged DNA depends on mono-ubiquitination of histone H2B (56). Notably, H2B ubiquitination was significantly diminished during infection with *S. pneumoniae* D39Δ*cps*, which was similar to the reduction of HERC4 and poly-ubiquitinated proteins observed before (Figure 6A and C and S4A and B, infection control, Figure 3A, 1C). In addition, both the decrease in ubH2B and the increase in pH2AX were reverted upon infection with *S. pneumoniae* D39Δ*cps*Δ*spxB* consistent with the findings for HERC4 and suggesting a role of HERC4 in DNA damage repair (Figure 6A - C, Figure 4C).

**Figure 6.**
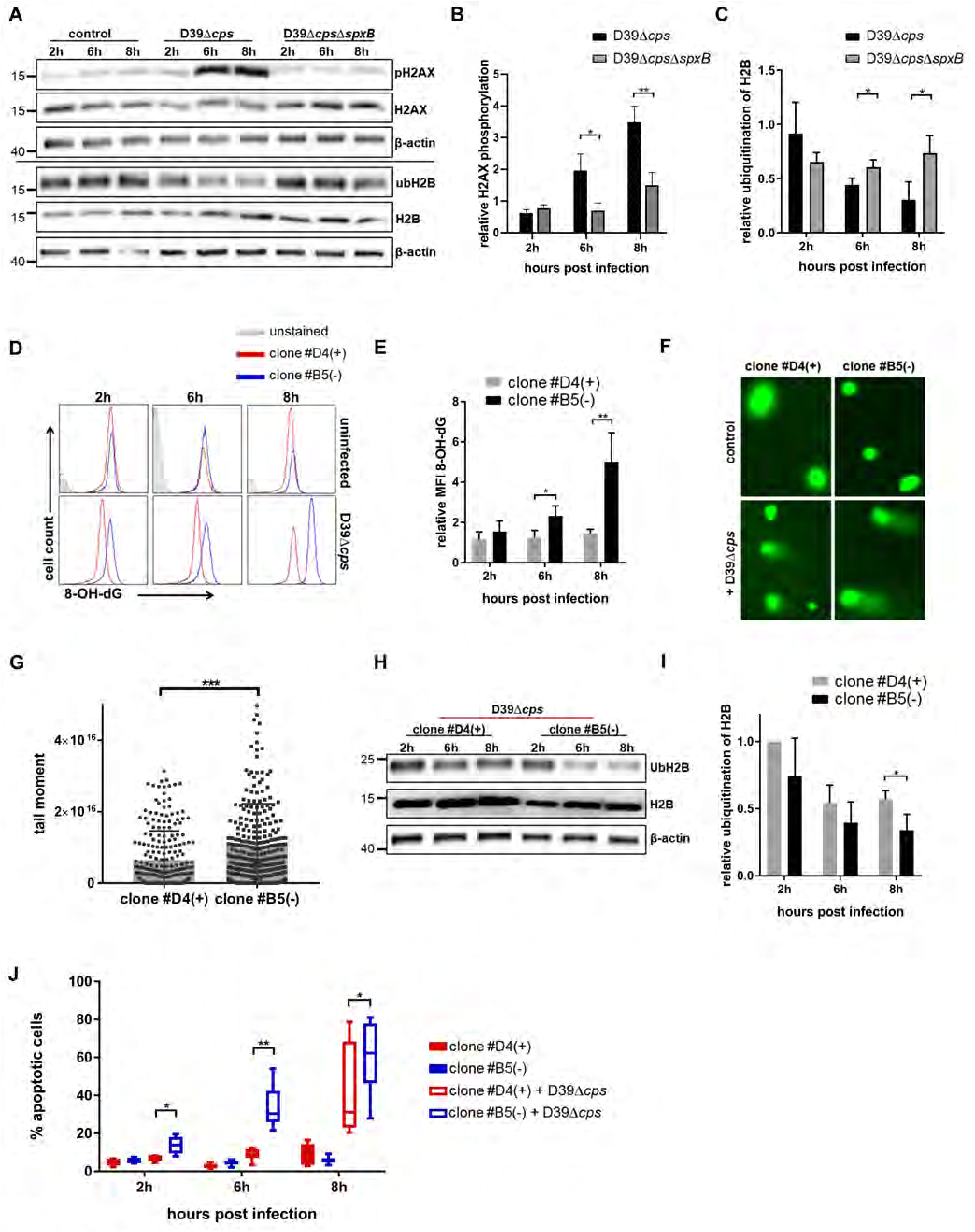
CRISPR-CAS9-mediated deletion of HERC4 fosters ROS-induced DNA damage and apoptosis. A–C A549 cells were analyzed for phosphorylation of H2AX and ubiquitination of H2B upon infection with *S. pneumoniae* D39Δ*cps* (MOI 20) and *S. pneumoniae* D39Δ*cps*Δ*spxB* (MOI20) compared to uninfected controls by immunoblotting (A). Band intensities of pH2AX were analyzed by densitometry and normalized to total H2AX and to the loading control β-actin. Graph depicts relative phosphorylation of H2AX calculated to 2 h uninfected control (B), n = 3. Band intensities of ubH2B were analyzed by densitometry and normalized to total H2B and to the loading control β-actin. Graph depicts relative ubiquitination of H2B calculated to 2 h uninfected control (C), n = 3. D-J CRISPR Cas9 mediated knockout of HERC4 in A549 cells, followed by infection with *S. pneumoniae* D39Δ*cps.* DNA damage was assessed by measuring the formation of 8-OH-deoxyguanosine by flow cytometry at indicated time points after infection in HERC4 KO cells (clone #B5) and control cells (clone #D4), MOI 5 (D) and calculated as fold induction of the MFI compared to the respective uninfected cells (E), n = 3. Direct DNA damage was assessed by COMET assay 8 h upon infection with *S. pneumoniae* D39Δ*cps* (MOI 5) (F), 240 calculated tail moments from three independent experiments were compared between infected HERC4 KO (clone #B5) and control cells (clone #D4) (G), n = 3. HERC4 KO (clone #B5) and control cells (clone #D4) were analyzed for ubiquitination of H2B during *S. pneumoniae* D39Δ*cps* infection (MOI 5) (H). Band intensities of ubH2B were analyzed by densitometry and normalized to total H2B and to the loading control β-actin. Graph depicts relative ubiquitination of H2B calculated to 2 h infected control cells (clone #D4) (I), n = 4. Annexin V / 7-AAD staining was used to detect the percentage of apoptotic cells in uninfected and *S. pneumoniae* D39Δ*cps* (MOI 5) infected HERC4 KO (clone #B5) and control cells (clone #D4), n = 6 (J). Data information: in B, C, E, I and J data are presented as mean ± SD. \**p* < 0.05; \*\**p* < 0.01; \*\*\**p* < 0.001. (student’s t test)

To further decipher the functional role of HERC4 during pneumococcal infections we generated A549 HERC4 CRISPR CAS9-knockout cell clones (Figure EV4A and B). These clones exhibited increased sensitivity to pneumococcal infections. Hence, a reduced MOI (MOI5) was required to obtain reliable kinetic data. In line with the data derived from wildtype A549 cells a significant increase in damaged DNA in the HERC4-deficient cell clones could be detected compared to infected HERC4-expressing control cells. This was verified by intracellular staining for 8’-hydroxy-2’-deoxyguanosine, a marker of oxidative DNA damage (Figure 6D and E) (57, 58). These results could be further confirmed by monitoring DNA damage in comet assays, showing a significant increase in the tail moments as a marker for DNA-fragmentation in infected HERC4 KO cells compared to infected HERC4-expressing control cells (Figure 6F and G).

Next, we addressed the question if HERC4 could have an effect on DNA repair processes. Upon infection with *S. pneumoniae* mono-ubiquitination of H2B was slightly decreased in control cells which was further reduced significantly in HERC4 knockout cells implicating an impaired ATM kinase recruitment to damaged DNA during infection (Figure 6H and I). Finally, analysis of cell viability upon infection revealed a significantly increased number of apoptotic cells in the HERC4-deficient cell clones #B5, #A8 and #C4 compared to control clone #D4, as determined by Annexin V staining (Figure 6J and EV4C - D and E, infection control). This was in line with the finding that the amount of oxidized proteins was further increased in HERC4 KO cells (Figure EV4F).

### HERC4 overexpression attenuates host cell DNA damage response during pneumococcal infections

The obtained results point towards a regulatory role of HERC4 during DNA damage repair. To determine whether increased HERC4 expression counteracts bacterial ROS-mediated host cell DNA damage, HeLa cells overexpressing HERC4 were generated by transfection with pCMV-Sport6-HERC4 and analyzed following infection with *S. pneumoniae* D39Δ*cps*. Compared to human alveolar epithelial cells such as A549 and primary SAEC cells, control transfected HeLa cells (pCMV-Sport6) displayed a similar extent of infection-induced HERC4 downregulation. In contrast, transfection with pCMV-Sport6-HERC4 attenuated HERC4-degradation, which was characterized by significant higher HERC4 levels 6 h post infection (Figure 7A and B, Figure S4C, infection control). In addition, elevated HERC4 mRNA levels were detected 24h after transfection (Figure 7C). As expected, increased HERC4 expression resulted in higher ubiquitination of H2B and reduced H2AX phosphorylation compared to control vector-transfected cells 6 h after infection pointing to decreased DNA damage and a diminished DNA damage response in pCMV-Sport6-HERC4 transfected cells (Figure 7D - F).

**Figure 7.**
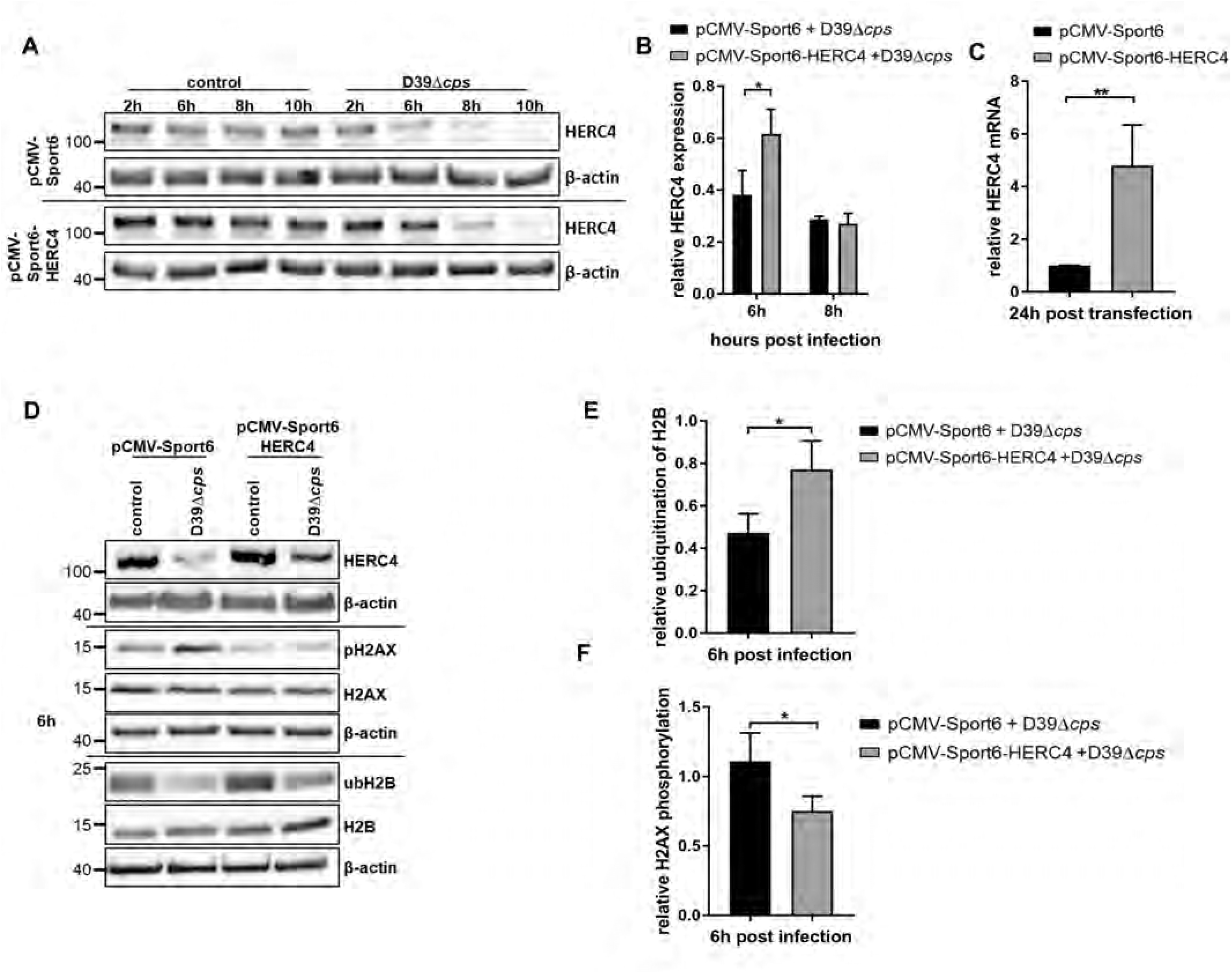
Decreased DNA damage upon S. pneumoniae infection of HeLa cells transfected with HERC4. A–C HeLa cells were transfected 24 h prior infection with pCMV-Sport6-HERC4 or the empty vector pCMV-Sport6 as control. Transfected cells were either infected with *S. pneumoniae* D39Δ*cps* (MOI 20) or non-infected (control) and analyzed at the depicted time points for HERC4 expression by immunoblotting (A). Graph depicts relative HERC4 expression at 6 h and 8 h calculated to 2 h uninfected control (B), n = 3. Changes in HERC4 mRNA were analyzed by qPCR 24 h after transfection. Relative HERC4 mRNA was calculated comparing HERC4 transfected and control transfected cells and normalized to the 18S rRNA housekeeping gene (C), n = 3. D–F pCMV-Sport6-HERC4 and pCMV-Sport6 transfected HeLa cells were infected for 6 h with *S. pneumoniae* D39Δ*cps* (MOI 20) and analyzed for HERC4, pH2AX, H2AX, ubH2B, H2B with β-actin as loading control (D). Graphs depict relative ubiquitination of H2B (E), n = 3, and the relative phosphorylation of H2AX (F), n = 3, at 6 h post infection in pCMV-Sport6-HERC4 and pCMV-Sport6 transfected cells compared to 6 h uninfected control, n = 3. Data information: in B, C, E and F data are presented as mean ± SD. \**p* < 0.05; \*\**p* < 0.01. (student’s test)

In summary, our findings uncover a link between pathogen-derived ROS, host cell ubiquitination and DNA damage during bacterial and viral-bacterial infection. Furthermore, we identified E3 ubiquitin ligase HERC4 as a potential regulator of host cell DNA damage response upon infection-induced oxidative stress.

## Discussion

In this study, we demonstrate that *S. pneumoniae* infection suppresses host cell ubiquitination, as evidenced by reduced levels of ubiquitinated proteins and downregulation of a defined subset of E2 and E3 ubiquitinating enzymes. Bacterial H_2_O_2_ contributes to, but does not fully account for this effect. The changes are associated with increased DNA damage, accumulation of carbonylated, oxidized proteins, and enhanced cellular toxicity. In contrast, infection with IAV H1N1pdm09 promotes the accumulation of ubiquitinated proteins and induces a distinct ubiquitination profile characterized by upregulation of ubiquitinating enzymes, which is consistent with an IFN-γ−driven signature observed in our previous work (36, 41) and in line with reports for other viral infections (30, 59).

Mechanistically, our data support a central role for bacterial H_2_O_2_ in modulating host cell ubiquitination. Notably, high concentrations of H_2_O_2_ (up to 1.83 mM) have been reported in culture supernatants of *S. pneumoniae* strains such as D39 (38, 60), representing levels that are unlikely to be reached intracellularly during viral infection. Accordingly, infection with the *S. pneumoniae* D39Δ*cps*Δ*spxB* mutant, which exhibits markedly reduced H_2_O_2_ production (61), resulted in diminished protein carbonylation to levels comparable to non-infected control cells and largely restored poly-ubiquitinated protein levels, as detected by an antibody that specifically recognizes poly-ubiquitinated proteins. These findings provide strong evidence that bacterial oxidative stress contributes to the observed changes in host cell protein ubiquitination. In addition to reduced expression of ubiquitinating enzymes during pneumococcal infections, high concentrations of H_2_O_2_ can block host protein synthesis via phosphorylation of the translation initiation factor eIF2α, thereby limiting the availability of newly synthesized proteins for ubiquitination (62, 63). At the same time, our data suggest that the regulation of individual E3 ligases cannot be explained by ROS alone but likely involves additional receptor-mediated signaling pathways.

ROS-generation in IAV-infected cells is assessed indirectly by quantifying the percentage of ROS-positive host cells rather than absolute concentrations (64), highlighting fundamental differences in the magnitude and measurement of oxidative stress between bacterial and viral infections. ROS generation in IAV-infected cells is strain-dependent. In line with this, infection with highly pathogenic avian H5N1 strains leads to reduced translocation of the antioxidant transcription factor nuclear factor erythroid 2-related factor 2 (NRF2) into the nucleus compared to seasonal and pandemic H1N1 viruses, suggesting an increased susceptibility of H5N1-infected cells to oxidative stress (65). Furthermore, IAV-induced ROS can sensitize host lung cells to bacterial toxin-mediated necroptosis during IAV – *S. pneumoniae* coinfection (66). Despite this, our data reveal that in coinfection settings, the ubiquitination profile is predominantly shaped by bacterial ROS, resulting in reduced expression of specific E3 ubiquitin ligases, such as HERC4, which is downregulated at both the mRNA and protein level but remains unchanged during IAV-monoinfection.

In contrast to *S. pneumoniae*, *S. aureus* (HG001 strain) mono-infection did not significantly alter the ubiquitinated protein levels or cellular protein carbonylation, consistent with its ability to detoxify hydrogen peroxide via catalase (67). However, during IAV – *S. aureus* coinfection, IAV-induced and host-derived ROS (particularly from NOX2-dependent pathways) have been shown to exacerbate oxidative stress and tissue damage, accompanied with an increased susceptibility to secondary *S. aureus* infection (68, 69). Together, these findings highlight that both pathogen - pathogen and pathogen - host interactions critically shape the cellular redox environment, with the nature and source of ROS emerging as critical determinants of downstream ubiquitination and cellular stress responses.

In our infection experiments with *S. pneumoniae*, we observed downregulation of a defined subset of ubiquitinating enzymes, including the E3 ligases HERC4 (and its interacting E2 enzyme UBE2L3, (70)), and LRSAM1. Mechanistically, downregulation of LRSAM1 was dependent on Toll-like receptor signaling, involving TLR2 and TLR4. In contrast, HERC4 expression was independent of Toll-like receptor signaling and instead linked to scavenger receptor (SR)-mediated pathways. Consistent with this finding, pulmonary surfactant protein A (SP-A)-mediated enhancement of *S. pneumoniae* phagocytosis by alveolar macrophages has been shown to depend on SP-A-induced cell surface localization of SR-A during infection (71). This suggests that scavenger receptors may contribute not only to bacterial uptake but also to the regulation of intracellular signaling pathways during *S. pneumoniae* infection. Notably, HERC4 levels were unaffected upon exposure to the H_2_O_2_-deficient *S. pneumoniae* D39Δ*cps*Δ*spxB* mutant but were reduced upon H_2_O_2_ treatment, suggesting a link between oxidative stress and scavenger receptor-dependent regulation of HERC4 during infection. Crosstalk between oxidative stress and scavenger receptor pathways has been reported primarily in the context of cardiovascular disease, including NRF2-dependent regulation of antioxidant proteins and scavenger receptor expression, as well as SR-A binding to oxidized proteins and lipids, which promotes inflammatory signaling in atherosclerosis (72). Together, these findings suggest that oxidative stress may engage scavenger receptor-dependent signaling, thereby contributing to pathogen-specific modulation of the host ubiquitination machinery.

In addition to the two E3 ligases described above, other E3 ligases showed a transient and less pronounced downregulation during *S. pneumoniae* infection. This effect may, at least in part, be explained by their auto-ubiquitination followed by degradation (73). In this context, oxidative modification of proteins such as E3 ligases and other (ubiquitinating) enzymes may promote their K48-linked ubiquitination and subsequent degradation by proteasomes, consistent with previous evidence linking protein oxidation to ubiquitin-dependent proteolysis (74). In our experimental system, E3 ligase HERC4 was downregulated at the transcript as well as the protein level, with the latter being degraded by both the proteasome and the lysosomal pathway. In contrast, in a previous study from our group, HERC4 was not found to be downregulated during IAV - *S. pneumoniae* coinfection, which may reflect differences in infection conditions, particularly the lower multiplicity of infection (MOI) used for *S. pneumoniae* in that study (41). Experimental knockout or overexpression of HERC4 further supported its role in cellular stress responses during *S. pneumoniae* infection. CRISPR-Cas9 knockout of HERC4 was associated with increased ROS-mediated DNA damage, induced apoptosis, and enhanced accumulation of oxidized proteins upon infection, whereas HERC4 overexpression attenuated these effects as shown by reduced phosphorylation of histone H2AX and increased mono-ubiquitination of histone H2B. *S. pneumoniae-*derived hydrogen peroxide has been shown to induce DNA damage in host cells, whereas host cell mechanisms act to attenuate this damage (50). In this context, E3 ubiquitin ligases have emerged as important regulators of DNA damage responses, controlling the stability and activity of key factors involved in genome stability (13, 75). Consistent with this, HECTD1 promotes base excision repair in chromatin through histone ubiquitylation, which stimulates the activity of apurinic/apyrimidinic endonuclease 1 (APE1) required for DNA damage repair after ionizing radiation (76). TRIM24 has been shown to be critical for DNA double-strand break (DSB)–induced recruitment of the Mre11–Rad50–Nbs1 (MRN) complex and activation of downstream ATM signaling in response to DSB induced by cancer therapy agents in human hepatocellular carcinoma cells, thereby facilitating efficient MRN-mediated DSB repair (77).

Supporting our findings in cellular infection models, analysis of patient-derived blood samples revealed differential expression of E3 ligase HERC4 in *S. pneumoniae -* versus IAV-infection, supporting the translational relevance of our findings. These observations were further underlined by *in vivo/ex vivo* data from *S. pneumoniae-*infected mice, in which HERC4 expression was downregulated in a strain-dependent manner, consistent with their differences in bacterial H_2_O_2_ production.

Thus, pathogen-specific regulation of host cell ubiquitination highlights ubiquitinating enzymes such as HERC4 as potential targets for therapeutic intervention in bacterial *S. pneumoniae* infection and IAV – *S. pneumoniae* co-infection. In addition, modulation of host cell ubiquitination may represent a strategy to limit pathogen-induced oxidative damage. Furthermore, defined alterations in the expression pattern of ubiquitinating enzymes identified here for IAV and *S. pneumoniae* may support the development of host-based diagnostic signatures to distinguish bacterial from viral infection and to identify patients at risk of bacterial co-infection during IAV-infection.

## Materials and Methods

### Cell lines

Human alveolar epithelial cell line A549, macrophage-like cell line THP1 and HeLa cells were cultivated in RPMI1640 (Gibco) medium containing 10% FCS (Capricorn). Human Small Airway epithelial cells (Lonza) were cultivated using Small Airway cell basal medium (SABM, Lonza) containing Small Airway Epithelial Cell Growth Medium BulletKit™ (SAGM, Lonza) as supplements.

### Bacterial strains

The *Streptococcus pneumoniae* serotype 2 D39Δ*cps* strain was used due to the fact that the interaction between the A549 cells and the pneumococci is mainly based on bacterial adherence to the cell surface which can be improved by using non-encapsulated strains (31, 32). D39Δ*cps*Δ*spxB* lacks also the bacterial pyruvate oxidase abolishing the release of hydrogen peroxide (38). Both strains were generated and described before (38) and maintained in complex medium Todd-Hewitt broth (Roth) supplemented with 0.5% yeast extract (THY) or on blood agar plates (BD Biosciences) at 37°C and 5% CO_2_. *S. aureus* HG001 was maintained in TSB (Roth) medium or on blood agar plates (BD Biosciences) at 37°C. For murine *in vivo S. pneumoniae* infection experiments, the serotype 4 strain TIGR4 (ATCC BAA-334)(78), a serotype 19F strain (BHN100)(79) and a serotype 7F strain (BHN54)(79) were used. Strains were obtained from B. Henriques-Normark (Karolinska Institutet, Stockholm, Sweden).

### Viral strains

Influenza A/H1N1/pdm09 was provided by U. Blohm (Federal research institute of animal health, Isle of Riems, Greifswald, Germany).

### Chemicals

The chemicals TLR2-IN-C29 (Selleckchem), TAK-242 (MedChemExpress), poly:I (Sigma Aldrich), bortezomib (Selleckchem), MG132 (Sigma-Aldrich), bafilomycin A1 (Selleckchem), and H_2_O_2_ (Carl Roth GmbH) were used for cell treatment throughout the study at the indicated concentrations and time points described.

### Bacterial, viral, and viral-bacterial infection experiments

For *S. pneumoniae* infections, cells were seeded into 6 well plates. 2 h before infection the medium of the cells was exchanged with RPMI without FCS. Bacteria were taken from a blood agar plate incubated at least for 7 h (37 °C, 5% CO_2_) and grown in Todd Hewitt broth supplemented with 0.5% yeast extract (THY) to mid-exponential phase (OD600 of 0.35 - 0.40). Pneumococci were washed and resuspended in PBS prior to infection. Cells were infected with *S. pneumoniae* D39Δ*cps* or D39Δ*cps*Δ*spxB* at a MOI of 20 unless otherwise indicated.

For viral infections, cells were infected at a MOI of 5. After 2 h the medium was replaced by RPMI supplemented with 10% FCS and cells were harvested at different time points post infection. For viral-bacterial co-infections, cells were infected with IAV with a MOI of 5 for 24 h followed by bacterial infection with *S. pneumoniae* D39Δ*cps* (MOI 20) for the depicted time points.

For *S. aureus* infections bacteria were taken from an overnight culture in TSB and grown in TSB media to mid-exponential phase (OD600 of 0.35 - 0.40), bacteria were washed and resuspended in RPMI media prior infection. Cells were infected at a MOI of 25. After 1 h infection 1 µg/µl Lysostaphin was added to prevent outgrowing of the bacteria.

For inhibitor experiments 8 h post infection serial dilutions from 10^-3^ – 10^-5^ of the cell culture supernatants were plated on blood agar plates. Colony forming units (CFU) were determined after 24 h incubation at 37 °C and the mean CFU numbers were calculated from all dilutions.

At the end of each infection period, cells were washed with PBS and detached using TRIzol^©^ reagent (Thermo Fisher Scientific).

### Murine pneumococcal infection experiments

Female C57BL/6JOlaHsd mice (age 10–14 weeks, purchased from Envigo) were used. Animals were housed in the animal facility of the Helmholtz Centre for Infection Research under SPF conditions and in accordance with national and institutional guidelines. The *in vivo* study protocol was reviewed and approved by institutional and regional ethical bodies (Niedersaechsisches Landesamt fuer Verbraucherschutz und Lebensmittelsicherheit, Nr. 33.19-42502-04-16/2319). Pneumococcal infection, isolation of primary type II alveolar epithelial cells (AECII) and microarray analyses were performed as previously described (80).

### CAPNETZ Study - setting and study population

Written informed consent was obtained from every patient or their legal representative before inclusion in the study. The study was approved by the local ethics committee (Nr. 301-2008, Hannover Medical School). Criteria for inclusion in this study were patients positive for *S. pneumoniae* in blood culture and patients positive for influenza A in respiratory samples. Disease parameters such as cough, production of purulent sputum, pathologic lung auscultation, fever, infiltrates, leukocyte number and the CRP were obtained through CAPNETZ (appendix table 1). For the present analysis whole blood samples of selected patients were analyzed for HERC4 expression via quantitative PCR.

### Immunoblot analysis

Proteins were isolated using TRIzol^©^ reagent (Thermo Fisher Scientific) according to manufactureŕs instructions and quantified via Bradford assay. For detection of oxidized proteins, the OxyBlot™ Protein Oxidation Detection Kit (Merck Millipore) was used according to manufacturer’s protocol. In brief 15 µg of cell lysate was derivatized by adding 10 µL of 1X DNPH solution for 15min. After adding the neutralization solution, the samples were ready to be analyzed by immunoblotting.

For immunoblotting proteins were separated by SDS-PAGE, transferred to nitrocellulose membranes, and analyzed for target proteins using primary antibodies listed in the reagent’s tools table. Membranes were developed with chemiluminescence using SignalFire™ ECL Reagent (Cell Signaling Technology) and analyzed with Imagequant 800 system (Cytiva). Densitometric quantification of the blots was performed by ImageJ (NIH).

### Quantitative PCR analysis

Total RNA from human cell lines was isolated using the Trizol reagent (Thermo Fisher Scientific) according to manufacturer’s instructions. Total RNA from patients infected with *S. pneumoniae* or influenza was isolated using Monarch total RNA Miniprep Kit (New England Biolabs). Reverse transcription was performed with M-MLV Reverse Transcriptase (Promega). Amount of HERC4 and NP mRNA was analyzed by qPCR. 18S RNA was used as housekeeping gene. The qPCR was performed using PowerUp™ SYBR™ Green Master Mix (Thermo Fisher Scientific). Used primers are listed in appendix table 2. For cell line infections relative expression levels of the selected transcripts were normalized to uninfected control cells and the housekeeping gene 18S rRNA and calculated using the 2^−ΔΔCt^ method. For patient samples relative expression levels of HERC4 were normalized to the housekeeping gene 18S rRNA and the 2^−ΔCt^ was calculated.

### LC-MS/MS sample preparation, analysis and data processing

After 10h *S. pneumoniae* infection proteins were recovered by resuspension of infected A549 cells in lysis buffer (50 mM Triethylammonium bicarbonate, 5% SDS) and DNA was sheared by sonication. The protein concentration was determined using the BCA assay (Pierce). Digestion of proteins was performed by suspension trapping on micro S Traps (Protifi) as described before (41). Briefly, lysates containing 20 µg protein were transferred into new tubes and disulfide bonds were reduced with TCEP followed by alkylation with iodoacetamide. Proteins were digested with trypsin (protease : protein - 1:25). Subsequently, peptides were eluted with 50 mM TEAB; 0.1% acetic acid; 60% acetonitrile in 0.1% acetic acid. Eluates were pooled and dried in a vacuum concentrator. Peptides were fractionated by basic reversed phase fractionation using the Pierce High pH reversed phase peptide fractionation kit (Pierce) according to the vendor protocol. Analysis was performed by LC MS/MS with an EASY nLC 1200 (Thermo Fisher Scientific) coupled to an Orbitrap Elite mass spectrometer (Thermo Fisher Scientific) operated in DDA mode. Detailed parameter can be found in appendix table 3. Raw data were searched with MaxQuant (version 1.6.0.16) (81, 82) against the UniProt databases for *Homo sapiens* and for *S. pneumoniae* D39. Identified protein groups were analyzed with Perseus (version 1.6.0.) (83, 84). After data filtering a two tailed t-test was applied for proteins with “LFQ intensity” values in three out of four replicates. Proteins were considered as differentially abundant with a p-value < 0.05 and with a fold change > 1.74 indicated as log2foldchange > |0.8|.

### Cytokine analysis

ELISA IL-8 Max Deluxe Set (BioLegend) was performed according to manufacturer’s instructions. In brief, plates were coated with capture antibody and incubated for 16-18 h at 4°C. Plates were washed (PBS + 0.5% Tween-20). Hereafter, plates were blocked with blocking solution for 1 h with shaking at 500 rpm. Furthermore, standard series were prepared (IL-8: 1000 pg/ml – 15.6 pg/ml). Non-specific bindings were blocked, blocking solution was removed and plates were washed. Standards and cell culture medium samples were added to the plates and incubated for 2 h with shaking. After the plates were washed, they were incubated with the detection antibody for 1 h. The detection antibody solution was removed, and plates were washed. Avidin-HRP solution was added and incubated for 30 min with shaking. Avidin-HRP solution was removed, and plates were washed. TMB substrate solution was added and incubated for 15 min in the dark. Stop solution (2 N H_2_SO_4_) was added and absorbance was measured at 450 nm and 570 nm within 15 min.

### Cell viability assays

The monitoring of cell viability was performed by analyzing the LDH content in cell supernatants by CytoTox-ONE Homogeneous Membrane Integrity Assay (Promega) according to manufacturer’s protocol. In brief 50 µL of supernatants were incubated for 10 min with 50 µL substrate solution. Afterwards the reaction was stopped by adding 25 µL of Stop solution. Supernatants of non-infected cells treated for 10 min at 37°C with 10 % TritonX 100 (AppliChem) served as positive control reflecting 100% cell death. Samples were measured at 590 nm using a plate reader (Tecan). To analyze apoptosis, cells were harvested using 1 % Trypsin (Capricorn Scientific) and subsequently stained with FITC Annexin V Apoptosis Detection Kit with 7-AAD (BioLegend). Data were acquired using a MACSQuant Analyzer 10 Flow Cytometer (Miltenyi Biotec) and analyzed with FlowJo™ Version 10.6.0 software.

### CRISPR-Cas9–mediated genome editing

CRISPR-Cas9 genomic editing for gene deletion was used as previously described (85). For *HERC4* gene deletion the sgRNA sequence 5’-GATCATCCGCAGATATCTCA-3’ was cloned into the pSpCas9 (BB)-2A-Puro plasmid (pX459, Addgene) and transfected into A549 cells. 48 h after transfection, cells were placed under puromycin selection (5 µg / mL) for 2 days and afterwards diluted to single cells. Clones recovered post single cell dilution were picked, grown, and identified by immunoblot analysis. Cell clones #B5, #A8 and #C4 were identified as HERC4 KO by immunoblotting whereas for clone #D4 no changes in HERC4 expression were observed and served as the control clone. Additionally, genomic DNA was purified and amplified by PCR using primers designed to flank the site targeted by the sgRNA. The PCR products were validated by Sanger sequencing (Eurofins Genomics).

### 8-hydroxy-2’-deoxyguanosine assay

Cells were harvested using 1 % Trypsin (Capricorn Scientific) at indicated time-points post infection. Afterwards, cells were fixed and permeabilized with Ebioscience FOXP3 intracellular staining Kit (Thermo Fisher Scientific) according to manufacturer’s instructions. Finally, cells were stained with 8-hydroxy-2-deoxyguanosine (8-OHdG)-FITC (Abcam, 1:100) and 7-AAD (BioLegend, 1:20). Data were acquired using a MACSQuant Analyzer 10 Flow Cytometer (Miltenyi Biotec) and analyzed with FlowJo™ Version 10.6.0 software.

### COMET assay

Single cell gel electrophoresis comet assays were performed using the SCGE assay Kit (Enzo Life Sciences). Following 8 h infection with *Streptococcus pneumoniae* D39Δ*cps* (MOI 5), cells were gently harvested with a cell scraper and mixed with low melting point agarose at a volume ratio of 1:50, and 75 μL aliquots were loaded onto pre-warmed slides. The slides were incubated in pre-chilled lysis solution for 45 min and then in pre-chilled alkaline solution for 45 min. Electrophoresis was run at 22 V in TBE buffer for 15 min. Comets were stained with CYGREEN dye for 30 min and imaged with fluorescent microscope REVOLVE (Discover Echo). At least 50 individual cells per sample were evaluated in duplicate using OpenComet Version 1.3 analysis tool.

### Transfection of HeLa cells

HeLa cells were transfected with 5 µg pCMV-Sport6-HERC4 vector (Mammalian Gene Collection, Dharmacon) or pCMV-Sport6 empty vector as control using ScreenFect A (ScreenFect) reagent 24 h prior infection.

### Statistical Analysis

Data were analyzed by comparing means using Student’s t-tests or multiple t-tests, as specified in the figure legends. All analyses were conducted with GraphPad Prism version 9.5.1. Results were considered statistically significant at *p < 0.05, **p < 0.01, and ***p < 0.001.

### The paper explained

#### Problem

Respiratory tract infections represent a major global health burden, with disease manifestation depending on the balance between pathogen virulence and host susceptibility. A pivotal driver of tissue damage in these infections is the accumulation of reactive oxygen species (ROS), which triggers a cascade of epithelial injury, genomic instability, and barrier dysfunction. This process is prominently observed in infections caused by *Streptococcus pneumoniae* and is further intensified during co-infections with Influenza A virus (IAV), which significantly exacerbates disease severity and patient outcomes. Despite the recognized role of oxidative stress in these infections, the intersection between pathogen-induced ROS and the host’s ubiquitin-dependent regulatory systems remains poorly understood.

#### Results

We demonstrate that *S. pneumoniae* produced ROS significantly reduces intracellular polyubiquitination and downregulates, among others, the E3 ligase HERC4. This downregulation occurs in human alveolar epithelial cells and macrophage-like cells, is mirrored in an infected mouse model and human patient samples, and is markedly intensified during IAV coinfection. Through CRISPR-Cas9-mediated deletion and overexpression, we show that HERC4 is essential for mitigating ROS-induced DNA damage, specifically by maintaining Histone 2B ubiquitination and facilitating DNA repair.

#### Impact

Our findings define the role of HERC4 as a guardian of genomic integrity during infection-induced respiratory stress. By uncovering a direct link between E3 ligase downregulation and impaired DNA-damage repair, this work provides a new mechanistic explanation for why *S. pneumoniae* infections and IAV – *S. pneumoniae* coinfections lead to severe cellular dysfunction. This identifies HERC4 as a critical determinant of infection outcome and a promising target for therapeutic intervention to attenuate the severe pulmonary pathology associated with respiratory pneumonia.

## Data availability

The mass spectrometry proteomics data have been deposited to the ProteomeXchange Consortium via the PRIDE (86) partner repository with the dataset identifier PXD079260. Reviewers can access the data via the PRIDE website using the following details: Project accession: PXD07926; Token: MCOsgWscog8.

Datasets analyzed for murine pneumococcal infection experiments are available in the Gene Expression Omnibus (GEO) repository under reference ID GSE225343.

## Acknowledgements

Research Training Group 2719 RTG-PRO, “Proteases in pathogen and host: importance in inflammation and infection” to SH and US, Mecklenburg-Western-Pomerania Excellence Initiative (Germany) and European Social Fund (ESF) Grant KoInfekt (ESF/14-BM-A55-0009/16) to UB, SH, DBe and US. This work was supported by the European Regional Development Fund (EFRE, 2014 – 2020; project number GHS-20-0031) to US and the Helmholtz Initialization and Networking Fund for Infection Research Greifswald (ZoonFlu; 2021) to DBru, UB, DBe and US. This study was supported by the German Research Foundation (TRR 359 - Project number 491676693 and BR2221/4 − 1, both to DBru).

We thank the patients for participation in this study. The investigators of this scientific work acknowledge CAPNETZ STIFTUNG and the CAPNETZ Study group for project support with regard to using biomaterials and clinical data (2023-03-07-GLX9_Seifert_Ubiquitin_E3-Ligases). CAPNETZ is a multidisciplinary approach to better understand and treat patients with CAP (https://capnetz.de/en/home/).

## Abbreviations

HERC4: HECT and RLD domain containing E3 ubiquitin protein ligase 4
ROS: reactive oxygen species
IAV: influenza a Virus
HECT: homologs to the E6-associated protein C-terminus LC-MS
LRSAM1: leucine rich repeat and sterile alpha motif containing 1
SAEC: small airway epithelial cells
TLR2: toll-like receptor 2
TLR4: toll-like receptor 4
SR: scavenger receptor
NOX2: NADPH Oxidase 2
NRF2: nuclear factor erythroid 2–related factor 2
UBE2L3: ubiquitin-conjugating enzyme E2 L3
ATM: ataxia-telangiectasia mutated protein kinase

## Author contributions

**Clemens Cammann:** Conceptualization; Methodology; Investigation; Formal analysis; Validation; Data curation; Visualization; Writing-original draft; Writing-review and editing. **Vanessa Gering:** Investigation; Methodology; Formal analysis. **Thomas Sura:** Investigation; Methodology; Formal analysis; Data curation; Writing-review and editing. **Abhishek K. Singh:** Investigation; Methodology; Formal analysis; Data curation; Writing-review and editing. **Julia D. Boehme:** Methodology; Investigation; Formal analysis; Validation; Writing-review and editing. **Eylin Topfstedt:** Investigation; Methodology; Formal analysis. **Anne K. Koch:** Investigation; Formal analysis. **Ulrike Ritter:** Investigation; Methodology; Formal analysis. **Karsten Becker:** Resources; Formal analysis; Writing-review and editing. **Dunja Bruder:** Resources; Methodology; Investigation; Formal analysis; Validation; Funding acquisition; Writing-review and editing. **Ulrike Blohm:** Resources; Methodology; Funding acquisition; Writing-review and editing. **Hortense Slevogt:** Methodology; Formal analysis; Writing-review and editing. **Sandra Maaß:** Investigation; Methodology; Formal analysis; Data curation; Writing-review and editing. **Gernot Rohde:** Resources; Formal analysis; Writing-review and editing. **Jan Rupp:** Resources; Formal analysis; Writing-review and editing. **Sven Hammerschmidt:** Resources; Methodology; Formal analysis; Validation; Supervision; Funding acquisition; Project administration; Writing-review and editing. **Dörte Becher:** Conceptualization; Resources; Methodology; Formal analysis; Validation; Supervision; Funding acquisition; Writing-review and editing. **Ulrike Seifert:** Conceptualization; Resources; Methodology; Formal analysis; Validation; Data curation; Visualization: Supervision; Funding acquisition; Project administration; Writing-original draft; Writing-review and editing.

## Expanded View Figure Legends

**Figure EV1.**
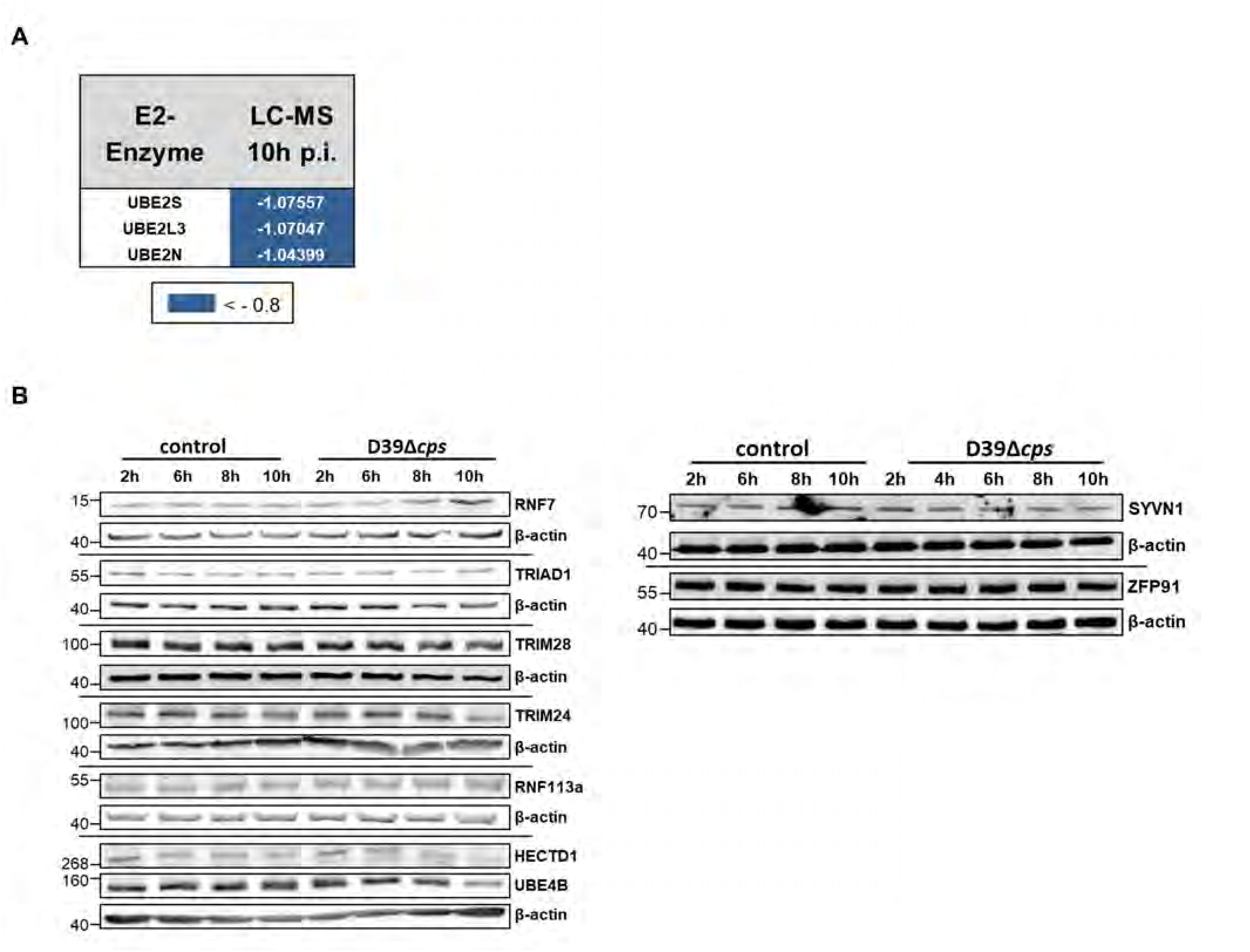
E3 Ligases during *S. pneumoniae* D39Δ*cps* infection. (Expanded view Figure related to Figure 2) A Comparison of differentially expressed E2 conjugating enzymes upon infection with *S. pneumoniae* D39Δ*cps* (MOI 20) which showed a significant decreased (blue) abundance in the LC-MS analysis after 10 h infection, n = 4. Proteins were considered as differentially abundant with a fold change > 1.74 indicated as log2foldchange > |0.8|, with p < 0.05. B A549 cells were infected with *S. pneumoniae* D39Δ*cps* (MOI 20) or non-infected (control) for the depicted time points and subsequently analyzed for E3 Ligase expression by immunoblotting with β-actin as loading control, one out of two experiments is shown, n = 2.

**Figure EV2.**
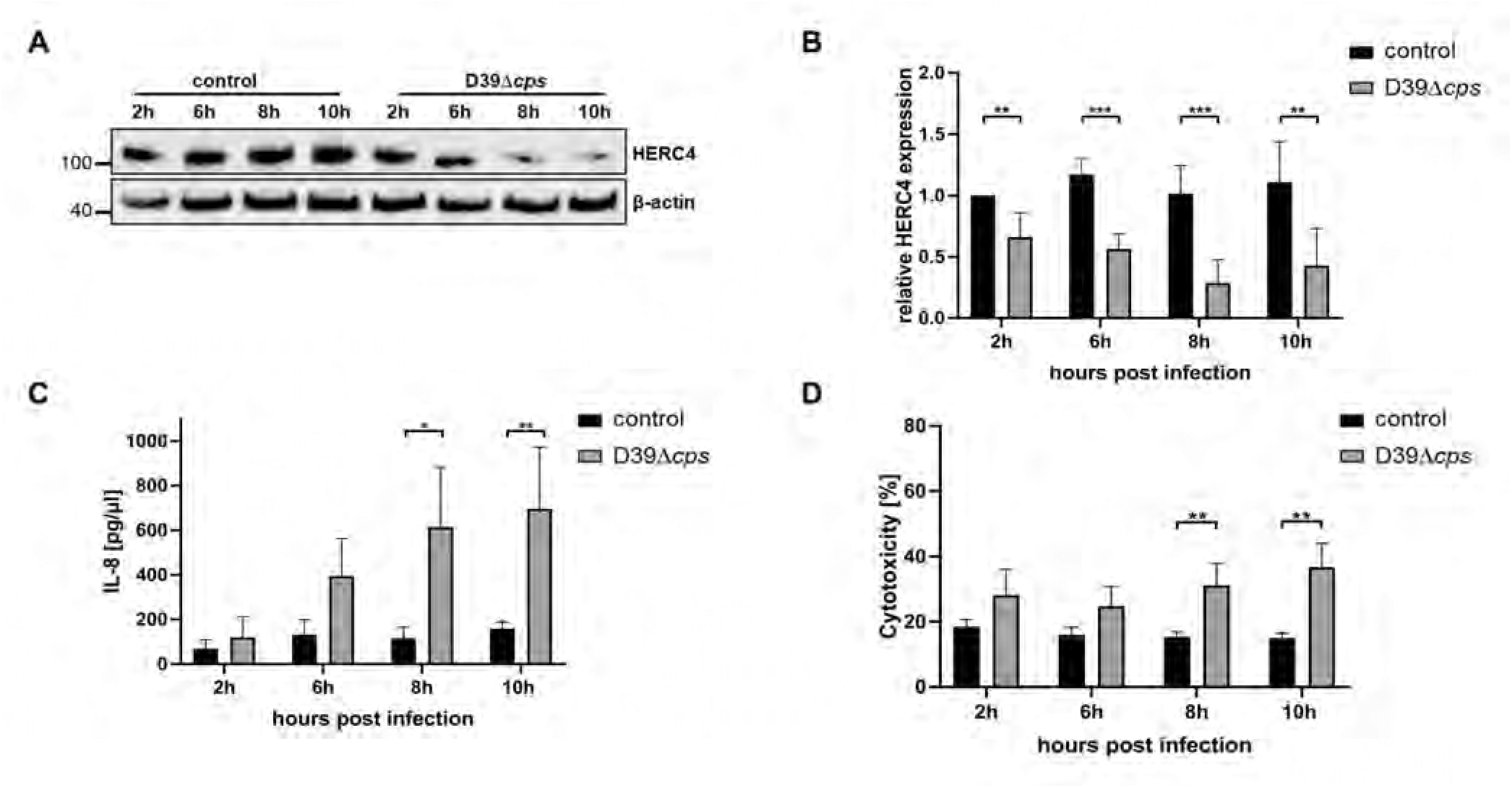
Reduced HERC4 expression in primary human small airway epithelial cells (SAEC). (Expanded View Figure related to Figure 3) A-D SAEC were infected with *S. pneumoniae* D39Δ*cps* (MOI 40) or left untreated (control) for the depicted time points and subsequently analyzed for HERC4 protein expression by immunoblotting with β-actin as loading control (A). Band intensities were normalized to the loading control β-actin. Graph depicts relative HERC4 expression calculated to 2 h uninfected control (B), n = 4. Infection controls: secretion of IL-8 was determined in the cell culture supernatants by ELISA (C), n = 4, and cytotoxicity was monitored by determining extracellular lactate dehydrogenase (D), n = 4. Data information: in B - D are presented as mean ± SD. *p < 0.05; **p < 0.01; ***p < 0.001. (B-D -multiple t tests)

**Figure EV3.**
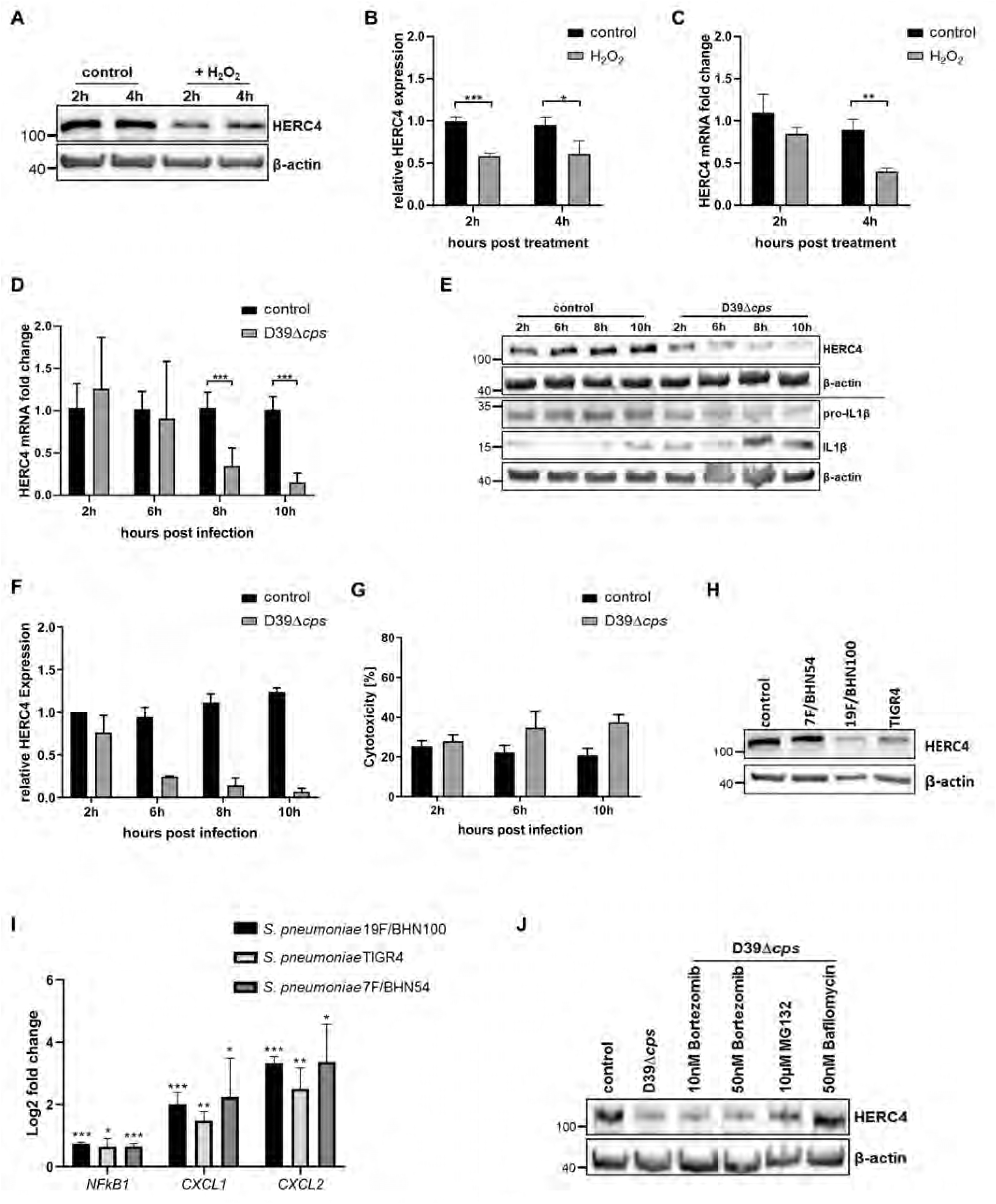
ROS induced downregulation of HERC4. (Expanded View Figure related to Figure 4) A, B A549 cells were treated with 500 µM hydrogen peroxide (H_2_O_2_) for the depicted time points or left untreated (control) and subsequently analyzed for HERC4 expression via immunoblot with β-actin serving as loading control (A), n = 3. Graph depicts relative change in HERC4 calculated to 2 h uninfected control (B), n = 3. C Changes in HERC4 mRNA upon H_2_O_2_ treatment was analyzed by qPCR, fold change was calculated to 2 h untreated control and normalized to the 18S rRNA housekeeping gene, n=3. D Changes in HERC4 mRNA upon *S. pneumoniae* D39Δ*cps* (MOI20) infection in A549 cells were analyzed by qPCR, fold change was calculated to 2 h uninfected control and normalized to the 18S rRNA housekeeping gene for *S. pneumoniae* D39Δ*cps* (D), n = 3. E-G THP-1 cells were differentiated for 72h with 100 ng / mL PMA. After 24 h resting they were infected with *S. pneumoniae* D39Δ*cps* (MOI 50) or left untreated (control) for the depicted time points and subsequently analyzed for HERC4, pro-IL1β and IL1β protein abundance by immunoblotting with β-actin as loading control (E), n = 2. HERC4 band intensities were normalized to the loading control β-actin. Graph depicts relative expression calculated to 2 h uninfected control (F), n = 2. Cytotoxicity was monitored by determining extracellular lactate dehydrogenase (G), n = 2. H Mice were oropharyngeally infected with 10^6^ *S. pneumoniae* (TIGR4, 7F/BHN54 or 19F/BHN100) or PBS (control). At 18h post infection type II alveolar epithelial cells were obtained from isolated lungs via fluorescence-activated cell sorting and pooled samples from 3-5 mice/experimental group were analyzed by immunoblotting for HERC4 expression with β-actin as loading control, n = 1. I Infection controls: Mice were oropharyngeally infected with 10^6^ *S. pneumoniae* (TIGR4, 7F/BHN54 or 19F/BHN100) or PBS. At 18h post infection gene transcription in type II alveolar epithelial cells was analyzed by microarray, *Nfkb1*/*Cxcl1*/*Cxcl2* genes were used as infection control calculated as log2foldchange and compared to PBS treated mice. Type II alveolar epithelial cells from n = 2-7 mice/experimental group/experiment were pooled. n = 3 J A549 cells were infected with *S. pneumoniae* D39Δ*cps* (MOI 20) and simultaneously treated with the depicted concentrations of bortezomib, MG132, and bafilomycin for 8 h and subsequently analyzed for HERC4 expression compared to untreated infected and non-infected cells (control) via immunoblot with β-actin serving as loading control, n = 2. Data information: in B-D and I data are presented as mean ± SD. *p < 0.05, **p < 0.01. ***p < 0.001. (student’s t test)

**Figure EV4.**
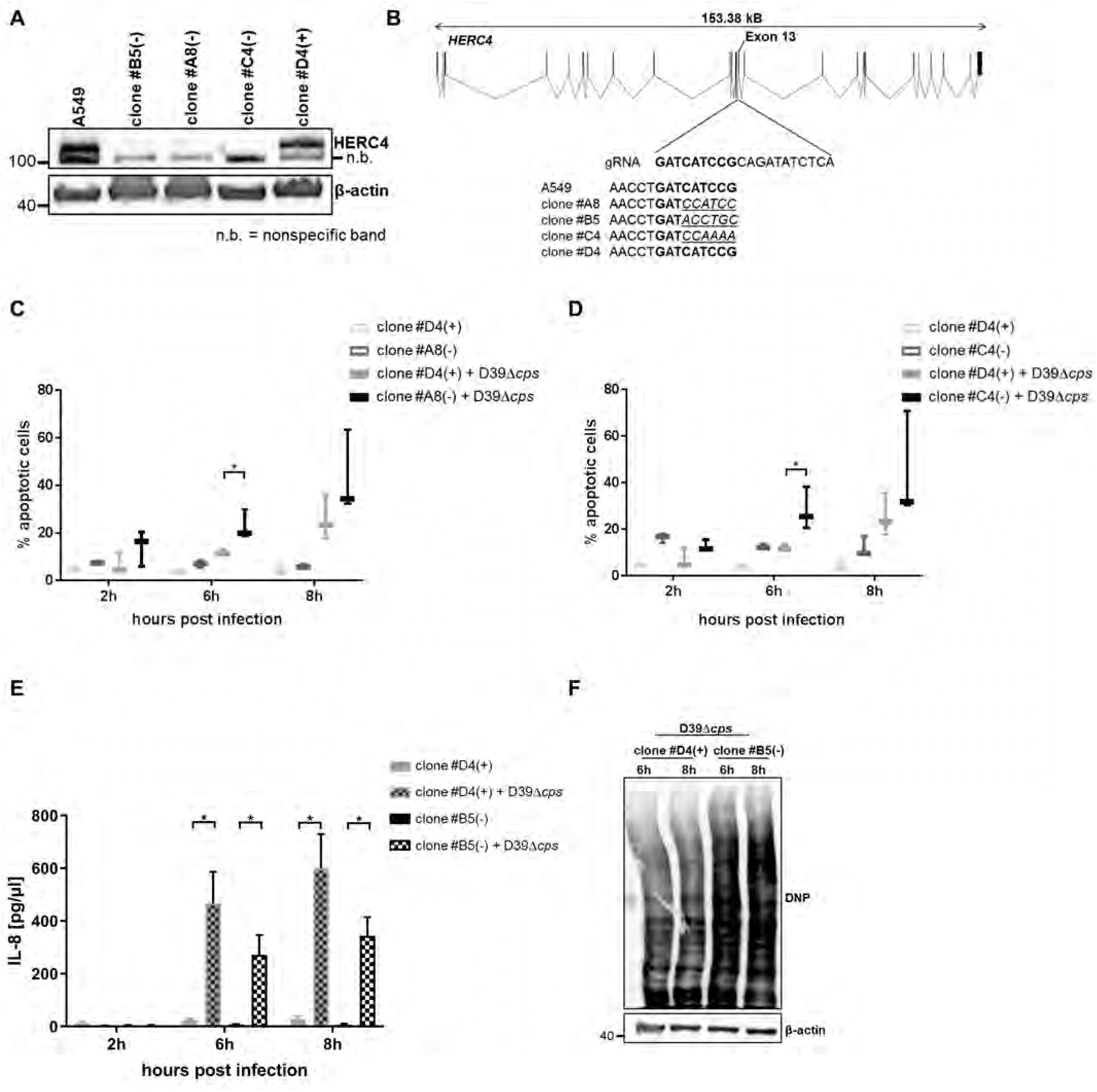
Deletion of HERC4 in A549 lung epithelial cells. A CRISPR CAS9 mediated deletion of HERC4 was confirmed by immunoblot for clone #A8, #B5 and #C4 and compared to control cells (A549 and clone #D4). B Sequencing of the genomic DNA of clones #A8, #B5 and #C4 confirmed CAS9-mediated cleavage at the designated site in exon 13 introduced by the gRNA sequence compared to the A549 parental cell line. Wild type sequences for A549 parental cell line and CRISPR Cas9 control cells (clone #D4) are printed in bold, missense sequences after CAS9 mediated cleavage are underlined in italics. C, D Annexin V / 7-AAD staining was used to detect the number of apoptotic cells at the depicted time-points in uninfected and *S. pneumoniae* D39Δ*cps* (MOI 5) infected HERC4 KO clone #A8 (D) and clone #C4 (E) and control clone #D4 (D and E), n = 3. E Secretion of IL-8 was determined for the depicted time points in the cell culture supernatants by ELISA during infection with *S. pneumoniae* D39Δ*cps* (MOI 5) of HERC4 KO (clone #B5) and control cells (clone #D4), n = 3. F HERC4 KO (clone #B5) and control cells (clone #D4) were analyzed for the amount of oxidized proteins via OxyBlot 6 h and 8 h upon *S. pneumoniae* D39Δ*cps* infection (MOI 20), n = 3, one representative blot out of three is shown. Data information: in C-E data are presented as mean ± SD. *p < 0.05. (student’s t test)

